# Interleukin-1 is overexpressed in injured muscles following spinal cord injury and promotes neurogenic heterotopic ossification

**DOI:** 10.1101/2021.10.19.464906

**Authors:** Hsu-Wen Tseng, Irina Kulina, Dorothée Girard, Jules Gueguen, Cedryck Vaquette, Marjorie Salga, Whitney Fleming, Beulah Jose, Susan M Millard, Allison R Pettit, Kate Schroder, Gethin Thomas, Lawrie Wheeler, François Genêt, Sébastien Banzet, Kylie A Alexander, Jean-Pierre Levesque

## Abstract

Neurogenic heterotopic ossifications (NHOs) form in periarticular muscles following severe spinal cord (SCI) and traumatic brain injuries. The pathogenesis of NHO is poorly understood with no effective preventive treatment. The only curative treatment remains surgical resection of pathological NHOs. In a mouse model of SCI-induced NHO that involves a transection of the spinal cord combined with a muscle injury, a differential gene expression analysis revealed that genes involved in inflammation such as interleukin-1β (IL-1β) were overexpressed in muscles developing NHO. Using mice knocked-out for the gene encoding IL-1 receptor (IL1R1) and neutralizing antibodies for IL-1α and IL-1β, we show that IL-1 signaling contributes to NHO development following SCI in mice. Interestingly, other proteins involved in inflammation that were also overexpressed in muscles developing NHO, such as colony-stimulating factor-1, tumor necrosis factor or C-C chemokine ligand-2 did not promote NHO development. Finally using NHO biopsies from SCI and TBI patients, we show that IL-1β is expressed by CD68^+^ macrophages. IL-1α and IL-1β produced by activated human monocytes promote calcium mineralization of fibro-adipogenic progenitors isolated from muscles surrounding NHOs. Altogether these data suggest that interleukin-1 promotes NHO development in both humans and mice.

## Introduction

Neurogenic heterotopic ossifications (NHOs) are pathological bone formations in soft tissues after central nervous system (CNS) injuries such as traumatic brain injuries (TBI), spinal cord injuries (SCI), strokes and cerebral anoxia. About 10-23% patients with TBI and 10-53% with SCI develop NHO and the prevalence increases to 68% in victims suffering from severe combat blast injuries^(1–3)^. NHOs develop in periarticular muscles and can lead to partial or complete ankylosis of affected joints and compression of blood vessels and nerves hence consequently compromises life quality of patients ^(4)^. Because our understanding of NHO pathogenesis is limited, the only effective curative treatment for NHO is surgical excision; however, even after surgical resection, NHO reoccur in 6-7% of patients ^(5)^. Preventive treatments such as radiation therapy and bisphosphonates have been proposed but may negatively impact other complications associated with SCI, such as wound and bone fracture healing ^(6)^. The only preventive treatments that show some efficacy at reducing the incidence and size of NHO in retrospective studies are non-steroid anti-inflammatory drugs (NSAIDs) such as indomethacin and celecoxib ^(7)^.

To better understand the cellular and molecular mechanisms driving NHO and discover potential therapeutic targets, we established the first SCI-induced NHO mouse model in which NHO development requires a combination of SCI and muscle injury without the artifice of genetic manipulation ^(8)^. In this mouse model, NHO development is induced by the combination of spinal cord transection between T11-T13 combined with a muscle injury induced by injection of cardiotoxin ^(8)^. In this model, NHOs are featured by mineralized calcium deposition detected by micro-computed tomography (μCT) from day 4 post-SCI and muscle injury. NHO volumes peak around day 7-14. As NHOs gradually mature, osterix-positive osteoblasts lining the NHOs lay down collagen I and osteocalcin-containing bone matrix and form small heterotopic bone nodules which are remodeled by multinucleated osteoclasts ^(8)^.

There are striking differences between normal muscle repair and NHO development. In the absence of SCI, muscle repair is completed within 3 weeks by satellite cells, which are myogenic stem cells that differentiate into myoblasts that fuse into myocytes to replace damaged myofibers. Effective muscle repair begins with acute inflammation in response to injury. CC chemokine ligand 2 (CCL2), also called macrophage chemoattractant protein 1 (MCP-1) and its receptor CC chemokine receptor 2 (CCR2), known to mediate monocyte recruitment in various inflammatory conditions, recruits inflammatory monocytes to the site of muscle injury ^(9–12)^. Inhibition of macrophage infiltration by depleting intramuscular CD11b^+^ myeloid cells, or deletion of CCL2 or CCR2 encoding genes significantly reduces infiltration of monocyte/macrophages and subsequently delays clearance of necrotic muscle and muscle regeneration ^(9,10,12,13)^. Recruited monocytes and muscle-resident macrophages produce cytokines that enhance myogenesis such as interleukin-6 (IL6), insulin-like growth factor 1 (IGF1) and interleukin-1β (IL1-β) ^(9,14)^. Another muscle-resident progenitor population called fibro-adipogenic progenitors (FAPs), expands rapidly following muscle injury and provides a supportive environment for satellite cell expansion and myogenic differentiation^(15)^. However unlike satellite cells, dividing FAPs undergo apoptosis in a tumor necrosis factor (TNF)-dependent manner between 4 and 7 days following injury and are eliminated by macrophages to prevent scar tissue formation and fibrosis ^(16)^. Therefore, muscle resident and infiltrating macrophages are pivotal to orchestrate several steps of muscle regeneration.

Interestingly, we and others have demonstrated that macrophages are also essential in HO formation in various mouse models ^(8,17,18)^. In our SCI-NHO model, we discovered abnormally increased inflammatory monocyte/macrophages infiltration in injured muscle 4 days following SCI and muscle injury ^(19)^. Depletion of macrophages significantly reduced the size of NHO, establishing the critical roles of macrophages in NHO ^(8)^. We showed that in the context of a SCI, macrophages infiltrating the injured muscle secrete abnormally high levels of oncostatin M (OSM) which drives NHO development ^(20)^. OSM exerts its effect in part by binding to a specific cell surface receptor made of GP130 complexed with the OSM receptor α chain (OSMR) which signals via JAK1 and JAK2 tyrosine kinases, which in turn tyrosine phosphorylate and activate the transcription factor STAT3 by promoting its translocation from the cytoplasm to the nucleus. JAK-STAT3 signaling is persistently activated at abnormally high levels in muscles developing NHO after SCI ^(19)^. Deletion of the *Osmr* gene or blocking OSM receptor downstream signaling with the selective JAK1-2 tyrosine kinase inhibitor ruxolitinib, significantly reduces NHO formation ^(19,21)^ demonstrating the important role of OSM/OSMR/JAK1-2 signaling in NHO development.

Since numerous macrophage-mediated inflammatory responses can promote muscle repair or fibrosis or ossification, our working hypothesis was that CNS injury skews these inflammatory responses toward fibrosis and ossification rather than muscle repair ^(22)^. In order to better understand which inflammatory pathways are activated following a SCI and how these promote NHO instead of muscle repair, we undertook gene expression profiling of muscles in the early phase following muscle injury. Gene set enrichment analysis (GSEA) showed that inflammatory responses, IL6-JAK-STAT3 and TNF-NFκB signaling gene sets were significantly enriched in muscles developing NHO with enhanced expression of OSM, colony stimulating factor (CSF)-1, TNF, CCL2 and IL-1β. Functional studies with selective inhibitors, neutralizing antibodies and knock-out mice revealed that among these pro-inflammatory cytokines, only OSM and IL-1 signaling positively contributed to NHO development after SCI. Finally, expression of IL-1β was confirmed in the muscular tissue surrounding NHO from patients with SCI or TBI and *in vitro* osteogenic assays of human FAPs isolated from muscles surrounding NHOs confirmed that both IL-1α and IL-1β produced by activated human monocytes promote calcium mineralization.

## Materials and Methods

### Animals

All experimental procedures were approved by the Health Sciences Animal Ethics Committee of The University of Queensland (# MRI-UQ/TRI/050/17) and followed the Australian Code of Practice for the Care and Use of Animals for Scientific Purposes. C57BL/6 mice were purchased from the Australia Resources Centre, B6.129S-*Tnfrsf1a*^tm1lmx^ *Tnfrsf1b* ^tm1lmx/J^ (abbreviated as *Tnfrsf1a/1b*^−/− −^), B6.129S4-*Ccr2^tm1Ifc^*/J (abbreviated as *Ccr*2^−/−^), B6.129S7-*Il1r1*^lm^/J (*Il1r1*^−/−^) mice were purchased from the Jackson Laboratory. *Il1a*^tm1Yim^ (*Il1a*^−/−^) and *Il1b*^tm1Yim^ (*Il1b*^−/−^) were kindly provided by Prof Yoichiro Iwakura ^(23)^ and backcrossed over into C57BL/6 background over 10 times. Experimental mice were maintained in specific pathogen-free animal facility 12-hour light/dark cycle with free access to standard chow pellets and water. Animals were randomly allocated to treatment groups and different genotypes were co-housed in same cages. Mice were identified with ear notches.

### NHO mouse model

For microarray analysis, six week-old female C57BL/6 mice underwent either spinal cord transection injury (SCI) at level T11-13 or sham surgeries followed by intramuscular (i.m.) injection of cardiotoxin (CDTX) purified from the venom of *Naja mossembica* snake (Sigma Aldrich) at 0.32mg/kg or saline into the hamstring muscle as previously described ^(8)^. Combinations of surgery and i.m. injections in each group (4 mice per group) were 1) SCI+CDTX, 2) SCI+saline, 3) Sham+CDTX and 4) Sham+saline. Two days post injury, mice were euthanatized by cervical dislocation and hamstring muscles were immediately harvested to extract total RNA for gene expression microarray analyses.

Samples for quantitative real-time reversed transcribed polymerase chain reaction (qRT-PCR) validation were collected from another cohort of mice two and four days post-injuries following identical surgical procedures with the following minor modifications ^(19)^ : 1) Purified CDTX from Naja pallida venom was from Latoxan, France (cat# L8102) due to discontinuation of N. mossembica CDTX production by Sigma-Aldrich, 2) saline was replaced by PBS, which does not cause muscle injury. All muscle samples were snap frozen and stored at −80 °C until RNA extraction.

For in vivo analysis of gene deletion and drug effects on NHO formation, mice underwent SCI surgery an injection of CDTX from Naja pallida as described above.

### RNA extraction

Whole muscle samples were homogenized in 4 ml of Trizol (Invitrogen) with TissueRuptorII (Qiagen Hilden) or IKA T10 basic homogenizer on ice for 30 seconds for 3 times with 10 second intervals followed by incubation at RT for 30 minutes. Tissue debris were removed by centrifuging at 12,000xg and supernatants were mixed with 1/5 volume of chloroform. After spinning down at 12,000xg, 4°C for 15 mins, aqueous phases containing RNA were transferred to fresh tubes for RNA precipitation with isopropanol. The RNA pellets were further washed in 75% ethanol, air-dried and dissolved in RNAse/DNase free water.

### Gene expression microarray analysis

RNA samples for microarray analysis were cleaned up using the RNeasy MinElute Cleanup Kit (QIAGEN) and the quantity and quality was verified using Bioanalyzer (Agilent Technologies, Inc.). Total RNA samples were amplified and purified using the Ambion Illumina RNA amplification kit (Ambion, Austin, USA) to yield biotinylated cRNA following the Illumina protocol, and further hybridized to MouseRef-8 v2.0 Gene Expression BeadChips and read on a IScan System using the IScan control software Version 3.5.31 (Illumina). Microarray data were analyzed by Illumina Genomestudio software (v2011.1), Gene Expression module (V1.9.0) using normalization algorithms with background subtraction followed by Illumina custom differential analysis algorithms. Normalized data are accessible on NCBI database with the assigned GEO accession number GSE165062. Genes with signal intensity higher than background (detection p value <0.01) in at least one sample were included for further analyses. Principle component analysis (PCA) was performed using R studio (Version 1.1.456) prcomp function while heatmap was generated using complexHeatmap packages^(24)^.

### Gene set enrichment analysis (GSEA)

Gene set enrichment analysis (GSEA; GSEA software v4.0.3) was performed using microarray signal intensity ^(25,26)^. In the present study, we selected the Hallmark gene set in the MSigDB molecular signature database v7.0. The Hallmark gene sets were generated by a computational methodology based on identifying overlaps between gene sets in other MSigDB collections including 50 well-defined biological processes that summarize and represent specific well-defined biological states or processes and display coherent expression^(27)^.

### qRT-PCR validation

For qRT-PCR validation, cDNA were synthesized from RNA samples using iScript cDNA kit (BioRad) per manufacturer’s instructions. mRNA expression was quantified using single-step qRT-PCR Taqman system. Reactions were prepared following instruction of TaqMan™ Universal PCR Master Mix using mouse *Il1b*, *Tnf*, *Osm*, *Ccl2*,*Csf1, Il1a* and ribosomal protein S20 (*Rps20)* (Supplementary Table 1). Ct values were normalized by the expression of house-keeping gene *Rps20*. The amplifications were performed using Applied Biosystems Viia7 Real-time PCR system.

### Micro computerized tomography (μCT) analyses

NHO volumes were measured in live mice in the Inveon PET-CT multimodality system (Siemens Medical Solutions Inc.) with parameters: 360° rotation, 180 projections, 80 kV voltage, 500 μA current, and effective pixel size 36μm. Mice were anaesthetized with 2% isoflurane oxygen mixture during imaging. After 3D image reconstruction, NHO volume was quantified in the Inveon Research Workplace (Siemens Medical Solutions Inc.) as previously described ^(19)^. To calculate NHO volumes, the region of interest (ROI) was drawn around the injured muscle containing calcification (excluding any normal skeletons). The quantification was not performed with any blinding, but after defining the ROI, the calcified regions were defined by setting the threshold Hounsfield units (HU) to 450 HU; therefore, only radiodensity > 450 HU was quantified as NHO. NHO development in *Ccr2*^−/−^ mice was repeated in 3 separate experiments. Due to instrument upgrade between experiments, the NHO formation in the first 2 experiments (WT, n=10; *Ccr2*^−/−^ n= 10) were measured in μCT scanner (μCT40, SCANCO Medical AG, Brüttisellen, Switzerland) at a resolution of 30 μm and three-dimensional (3D) images of the lower parts of the mouse bodies were reconstructed from the scans using the μCT system software package. Quantitative assessments of bone volumes in the muscular mass by a subtraction technique of orthotopic mouse skeleton (hip, femur, tibia, fibula) were measured to detect and quantify NHO. The remaining *Ccr2*^−/−^ and wild-type mice (WT, n=5; *Ccr2*^−/−^ n= 3) were measured in the Inveon PET-CT scanner. All other mice were analyzed in the Inveon PET-CT scanner.

### In vivo treatment with GW2580 and anti-IL1α/anti-Ilβ neutralizing antibodies

C67BL/6 mice were administered with GW2580 (80mg/kg in 0.5% hydroxypropyl methylcellulose in water, twice daily oral gavage)^(28)^ or vehicle from day 0 to day 10 post-surgery. To confirm the effects of GW2580 on myeloid cell development, bone marrow (BM) and blood samples were analyzed by flow cytometry.

C67BL/6 mice were also injected intraperitoneally with 200 μg daily of neutralizing Armenian hamster anti-mouse IL-1α (clone ALF-16) together with 200 μg neutralizing Armenian hamster anti-mouse IL-1β (clone B122) monoclonal antibodies or 200 μg control Armenian hamster IgG monoclonal antibody (clone PIP) from day 0 to day 7 post surgery.

### Phenotypic characterization of mouse leukocytes by flow cytometry

Muscle leukocytes were isolated from hamstring muscles four days post-surgery using a skeletal muscle dissociation kit (Miltenyi Biotech, cat#130-098-305) and GentleMACS Dissociator tissue homogenizer (Miltenyi Biotec) as previously described ^(19)^. To characterize inflammatory infiltrating injured muscle in C57BL/6 and *Ccr2*^−/−^ mice, leukocytes were stained with the following cell surface markers: anti-lineage (B220, Ter119, CD3∊)-FITC, anti-Ly6C-Pacific blue, anti-F4/80-APC, anti-LY6G-PECy7, CD48-APC-CY7, CD11b-BV510, CD45-BV785 and anti-CCR2-PE (Supplementary Table 2). Stained cells were analyzed with CytoFlex flow cytometer (Beckman Coulter) equipped with 405 nm, 488nm, 561 nm and 635 nm lasers.

To confirm effect of GW2580 on myelopoiesis, separate cohorts of mice were injected subcutaneously with either saline control 1 mg/kg/day long-lived porcine CSF-1-IgG Fc fusion protein that cross-reacts with mouse as previously described ^(29)^. These two cohorts were both subdivided into two groups and gavaged with either vehicle control or 80 mg/kg GW2580 twice daily. Mice were treated for two consecutive days and femoral bone marrow (BM) harvested the next morning. BM from one femur was flushed with 1mL PBS containing 2% newborn calf serum (NCS) in a 1 mL syringe mounted with a 25G needle and washed in PBS/2% NCS prior staining. BM cells were subsequently stained with CD11b-BV605, anti-F4/80-AF647, anti-Ly6G-PECY7, CD115-PE, anti-Ly6C-Pacific Blue and CD48-PercPCY5.5 monoclonal antibodies (Supplementary table 2). Cells were analyzed using a CyAn flow cytometer (Beckman Coulter) equipped with 405 nm, 488nm and 635 nm lasers. All FCS data files were exported without compensation and analyzed on FlowJo v10.7.1 software following compensation with single color controls.

### Human ethics statement and samples collection

Blood, NHO and muscle samples were obtained with the informed consent of the patients and under the approval of an independent ethics committee (CPP approval n°09025, study BANKHO). Blood samples from Healthy donors (HD) and NHO patients were collected on citrate solution from Percy hospital (Clamart, France) and Raymond Poincaré hospital (Garches, France). NHOs and muscle surrounding NHOs were collected from surgical waste following their excision from patients with brain injuries (TBI) or spinal cord injuries (SCI) at Raymond Poincaré Hospital (Garches, France).

### Human CD68 and IL-1β immunohistochemistry

Human NHO biopsies were fixed in 4% formaldehyde, demineralized in 10% nitric acid and embedded in paraffin. 4 μm sections were deparaffined in xylene and rehydrated. For IHC staining, sections were permeabilized 15 min at room temperature (RT) in PBS-0.1% Triton X-100 followed by a 15 min treatment at 95°C in an pH=6.0 antigen retrieval citrate-based solution (Vector Laboratories, Burlingame, CA, USA) then cooled down for 15 min. Sections were then incubated in Bloxall blocking solution (Vector Laboratories) for 10 min at RT to inactivate endogenous peroxidase followed by a 30 min incubation in PBS-GS 10% for 30 min at RT. Anti-human CD68 (mouse monoclonal IgG3, clone PG-M1, Dako, dilution 1/100) and anti-human IL-1β (rabbit polyclonal antibody, NB600-633, Novus Biologicals, dilution 1/200) primary antibodies were diluted in PBS containing 5% goat serum and 1.5% bovine serum albumin (BSA) and incubated overnight at 4°C. Mouse IgG3 isotype (MAB007, R&D systems) and rabbit polyclonal isotype (ab37415, Abcam) were used as controls. After washing in PBS-Triton 0.1%, ready to use ImPRESS FLEX REAGENT goat anti-rabbit IgG and goat anti-mouse IgG (Vector Laboratories) were incubated for 30 min at RT. After washing in PBS-Triton 0.1%, FLEX DAB substrate solution (Vector Laboratories) was added on sections for 5 min at RT. Counter coloration was performed using hematoxylin. All sections were analyzed using a Panoramic Midi II slide scanner and Case Viewer software (3D HISTECH Ltd.).

### Human plasma IL-1β quantification by ELISA

Blood samples were centrifuged at 2,000 rpm for 10 min at 4°C and plasma fractions were stored at −80°C. IL-1β plasma level was evaluated using Human IL1-β ELISA Kit (DLB50, R&D Systems) according to the manufacturer procedure.

### Isolation of PDGFRα^+^CD56^-^ fibro-adipoprogenitors (FAPs) from human muscles in surgically resected NHO

Muscle fragments associated with NHO biopsies were minced using scalpel and small scissors, placed in a 50 ml Falcon tube and incubated in 1.5 mg/ml pronase (cat#10165921001, Sigma-Aldrich) in α-minimal essential medium (α-MEM), 45min in a 37°C water bath. After addition of α-MEM supplemented with 15% FBS and 1% P/S antibiotics, the cell suspension was filtered through a 100 μm cell strainer followed by a 40 μm cell strainer (BD Falcon). Isolated muscle progenitor cells (MPCs) were maintained 10 days in α-MEM supplemented with 15% FBS, 1% P/S antibiotics and 10 ng/ml basic fibroblast growth factor (bFGF) (R&D Systems). Human MPCs were trypsinized and incubated 30 min with biotinylated anti-human PDGFRα (BAF322 R&D Systems) goat polyclonal antibody and CD56-PE (clone B159, BD Pharmigen) monoclonal antibody in PBS 2% fetal bovine serum (FBS), 2mM EDTA or with control isotypes IgG1 PE (A07796, Beckman Coulter) and biotinylated goat IgG (BAF108, R&D Systems). Cells were washed and incubated 30 min with Streptavidin-APC/Cy7 and the viability dye 7-aminoactinomycin D (7-AAD) (Sony). Cells were washed and filtered through a 30 μm cell strainer (Sysmex) and sorted using a FACSAria III SORP sorter (BD Biosciences). PDGFRα+CD56-FAPs were seeded at 3,000 cells per cm2 in α-MEM supplemented with 10% FBS and 1% P/S antibiotics.

### Human CD14^+^ monocytes isolation and preparation of conditioned medium (CM)

Human CD14+ blood monocytes were isolated from whole blood donation buffy coat from the Centre de Transfusion Sanguine des Armées (Clamart, France) or NHO bone marrow. PBMC and NHO-bone marrow mononuclear cells were recovered from using Ficoll gradient (Pancoll, Pan-Biotech). CD14+ monocytes were then isolated following magnetic separation using CD14 microbeads (130-050-201, Miltenyi Biotec). CD14^+^ cells were seeded at 1.25×10^6^ cells per cm^2^ and stimulated with 100 ng/mL LPS (L6529, Sigma-Aldrich) or without LPS for 72h. Conditioned media with (CM^+^) or without (CM^-^) LPS stimulation were collected, centrifuged 5 min at 500g. Pooled CM^+^ were used in a 1 to 10 ratio in cell culture medium during osteogenic differentiation assays. Due to a limited number of CD14^+^ monocytes retrieved from NHO bone marrows, only CM^+^ derived from CD14^+^ blood monocytes were used. IL-1β and IL-1α concentrations in conditioned media were evaluated using Human IL1-β and IL-1α ELISA assays (DY201 and DY200, R&D Systems) according to the manufacturer’s instructions.

### *In vitro* osteogenic differentiation assay and mineralization quantification

Human PDGFRα+ CD56-FAPs were seeded at 3,000 cells per cm2 in α-MEM 10% FBS with standard penicillin / streptomycin (P/S). After 3 days, culture medium was replaced by control medium (α-MEM 10% FBS 1% P/S antibiotics) or osteogenic medium (α-MEM 10% FBS 1% P/S antibiotics, 0.052 mg/mL dexamethasone, 12.8 μg/mL ascorbic acid and 2.15 mg/mL β-glycerophosphate (D-1159, A-8960, G-9422, Sigma-Aldrich) for 13 days. Human recombinant IL-1α, IL-1β and IL-1RA were purchased from PeproTech (200-01A, 200-01B and 200-01RA). FAPs were incubated 2 hours with IL-1RA prior to conditioned medium (CM) addition. Neutralizing mouse anti-human IL-1α and human IL-1β monoclonal IgG1 antibodies and mouse IgG1 isotype control (mabg-hil1a, mabg-hil1b, mabg1-ctrlm) were purchased from InvivoGen and used at the indicated concentration with a 2 h pre-incubation in CM+ prior to addition to FAPs. At the end of the differentiation process, cells were washed in PBS, fixed in 70% ethanol, washed in water, and stained in 20 g/L Alizarin red S (A5533, Sigma-Aldrich). Excess stain was removed during 3 washing steps. Alizarin Red S dye was extracted with 0.5N hydrochloric acid and 5% sodium dodecyl sulfate (SDS) and quantified by spectrophotometry at 405 nm.

### Western blot

Cultured human PDGFRα+CD56-FAPs were washed in PBS prior to incubation in cell lysis buffer composed of PBS, 1% NP-40, 1% Triton X-100, 0,1% sodium dodecyl sulfate (SDS), 0,5% deoxycholic acid and protease inhibitors (11697498001, Roche). Cells were then scrapped, incubated 20 min on ice and supernatant was collected after centrifugation (13 000g, 15min at 4°C). Protein concentrations were quantified using Pierce BCA protein assay kit following manufacturer instructions (23252, ThermoFisher). 20 μg of protein were loaded for separation in 10% acrylamide SDS-PAGE gels (161-0182, Biorad) before transfer on PVDF membrane (Trans-Blot Turbo, Biorad). After saturation in 5% non-fat dry milk, membranes were incubated with specific RUNX2 antibody (dilution 1/1000, ab192256, Abcam) overnight at 4°C. Then, they were probed with HRP secondary antibody (dilution 1/200 000, ab205718, Abcam) 2h at RT. Following enhanced chemoluminescence incubation (170-5061, Biorad), imaging and densitometric analysis were performed on a ChemiDoc XRS+ (Biorad) using ImageLab for StainFree normalization.

### Statistical analyses

Statistical analyses were performed using non-parametric two-sided Mann-Whitney test for 2 comparisons or repeated measures Friedman test with Dunn’s correction or ANOVA with Tukey’s post-hoc test for multiple comparisons depending on normality of data distribution as specified in figures.

## Results

### SCI exacerbates inflammation in injured muscles as determined by expression microarray analyses

We first performed differential gene expression microarray analyses in mouse muscles 2 days following SCI and muscle injury to identify potential triggers of NHO development in response to SCI. To this end, a gene expression microarray analysis was performed using total RNA according to following four treatment groups: *1)* SCI+CDTX: SCI surgery plus intramuscular (i.m.) CDTX injection, the only condition in which NHOs form in the injured muscle; *2)* SCI+Saline: SCI surgery plus i.m. injection of saline; 3) Sham+CDTX: Sham surgery plus i.m. injection of CDTX; or *4)* Sham+Saline: Sham surgery plus i.m. injection of saline. Normalized data are accessible on NCBI database with the assigned GEO accession number GSE165062.

We noticed that one SCI+CDTX sample showed similar gene expression signatures to that of Sham+CDTX group (not shown). Incomplete spinal cord transection can occur during surgery, however, it takes more than two days to observe motor recovery. Therefore, we were unable to confirm if the spinal cord transection injury was complete or not in this particular mouse by day 2. As a result, this sample was excluded from further analysis.

Principal component analysis (PCA) of the remaining samples demonstrated that the dominant principal component 1 (PC1) segregated samples according to CDTX-mediated muscle injury while PC2 segregated samples according to SCI (Fig. 1A). As a result, all muscle samples specifically clustered according to treatment groups.

**Fig 1.**
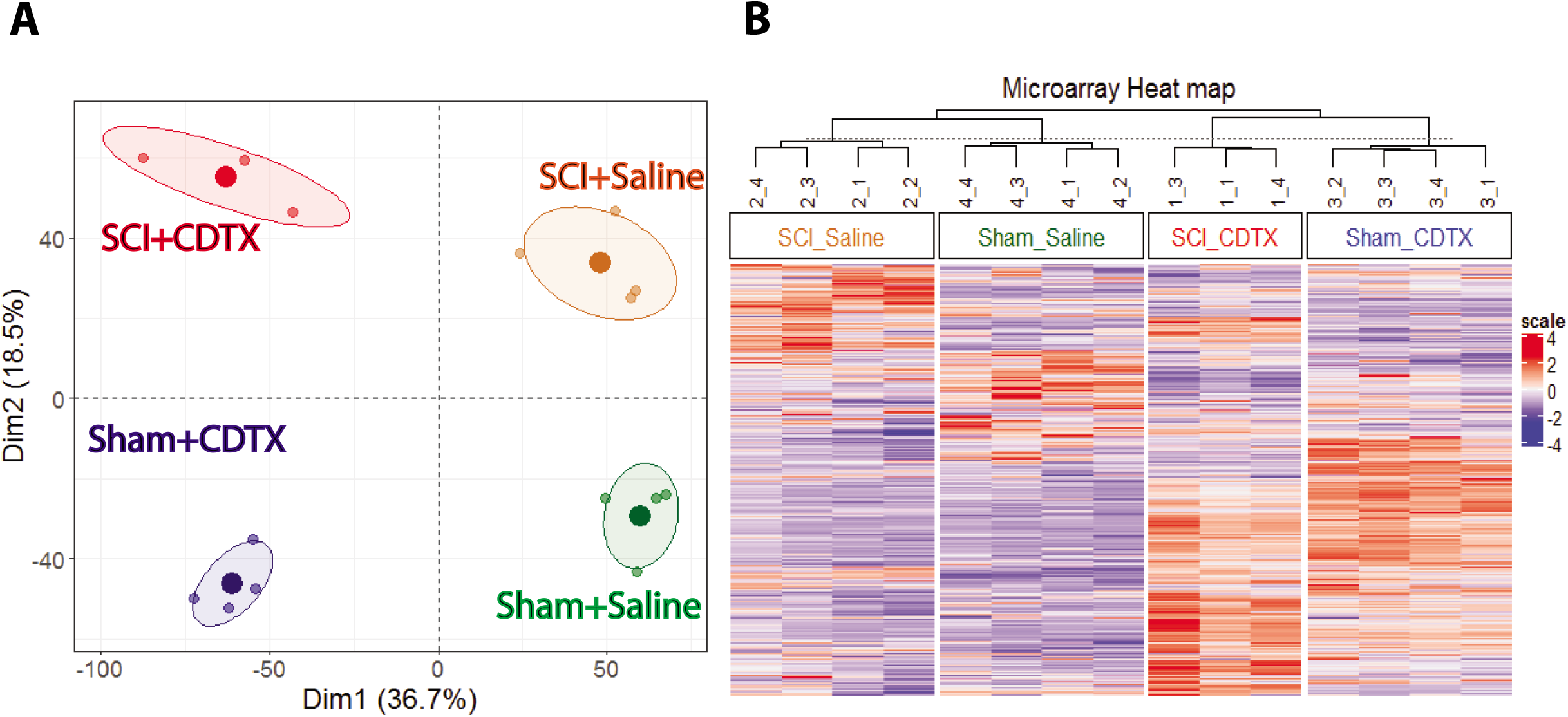
SCI causes expression of different sets of genes in injured skeletal muscles. RNA was isolated from injured hamstring muscles two days post-injury from four experimental groups: *1)* SCI+CDTX (red): SCI plus intramuscular (i.m.) CDTX injection, the only condition in which NHOs form in the injured muscle; *2)* SCI+Saline (orange) : SCI surgery plus i.m. injection of saline); *3)* Sham+CDTX (blue): Sham surgery plus i.m. injection of CDTX, ; or *4)* Sham+Saline (green): Sham surgery plus i.m. saline injection of saline. (A) principal component analysis (PCA) analysis. Each dot represents an individual mouse. (B) Clustering analysis with dendrogram and heatmap of differentially expressed genes in the four groups.

Expression heatmap confirmed that the combination of SCI plus CDTX muscle injury regulates unique sets of genes in the injured muscle that can be distinguished from genes activated in response to the SCI or CDTX injury alone (Fig. 1b).

GSEA were subsequently performed to identify which pathways and genes may be important in triggering NHO development by comparing differentially expressed gene sets between pairs of treatment groups using MSigDB Hallmark gene set collection ^(25–27)^. We performed the following comparisons:

Comparison 1: SCI+CDTX group vs the other 3 experimental groups (other) (Fig. 2A)
Comparison 2: SCI+CDTX group vs Sham+CDTX group (Fig. 2B)
Comparison 3: SCI+saline group vs Sham+Saline group (Fig. 2C)
Comparison 4: SCI+CDTX group vs SCI+Saline group (Fig. 2D)
Comparison 5: Sham+CDTX group vs Sham+Saline group (Fig. 2E)

**Fig 2.**
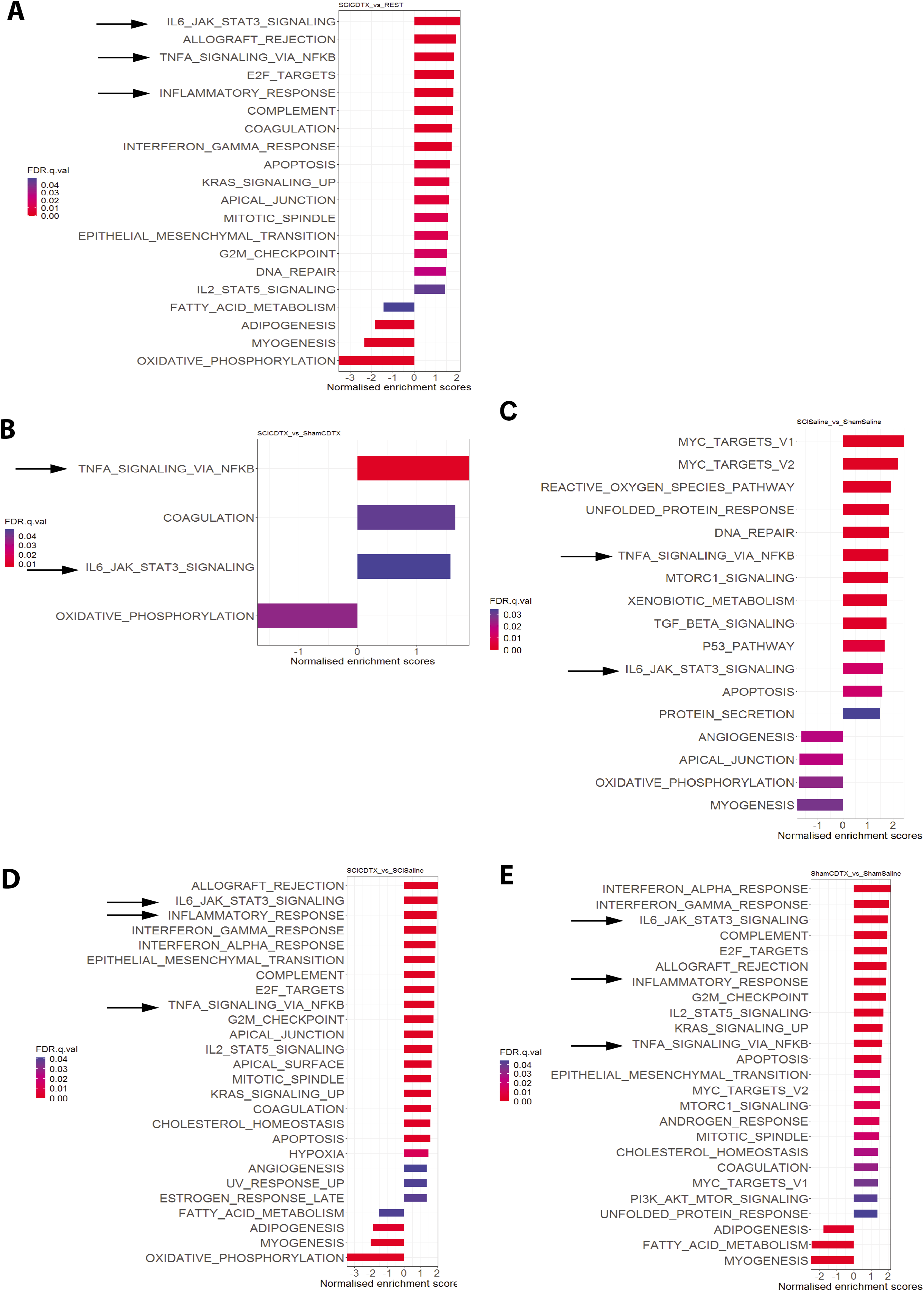
Gene set enrichment analyses of muscles from mice with SCI and /or muscle injury. Gene set enrichment analysis was performed to investigate genes enriched in injured muscles from the following experimental groups: (A) SCI + CDTX group which is the only developing NHO versus the other 3 experimental groups; (B) SCI + CDTX group versus Sham + CDTX group; (C) SCI + saline group versus Sham + Saline group; (D) SCI +CDTX group versus SCI + Saline group; (E) Sham + CDTX group versus Sham + Saline group. Gene sets with false discovery rate (FDR) below 0.05 are shown in the figures. Normalized enrichment scores (NES) were presented in X axes and FDR are represented on a color scale from high (blue) to low (red). Black arrows highlight the “inflammatory response”, “IL6-JAK-STAT3 signaling” and TNF signaling via NFκB” gene sets across the different comparisons.

Analysis of normalized enrichment scores (NES) with significant false discovery rates (FDR q <0.05) revealed that seven out of the eight top gene sets enriched in the SCI+CDTX group developing NHO relative to the 3 other treatment groups that do not develop NHO (Fig. 2A), were gene sets related to inflammation such as “IL6 JAK-STAT3”, “Allograft rejection”, “TNFA signaling via NFKB”, “inflammatory response”, “Complement”, “Coagulation”, “Interferon gamma response” further demonstrating that SCI exacerbates the inflammatory response in injured muscles as early as day 2 post-injury. Conversely, the three gene sets that were most significantly reduced in the SCI + CDTX group were “oxidative phosphorylation”, “myogenesis” and “fatty acid metabolism” suggesting that muscle repair and muscle energy usage are reduced subsequent to a SCI. Enriched gene sets in both comparisons 2 and 3 were more likely to be the consequence of the SCI. Enriched gene sets in both comparisons 4 and 5 revealed gene sets enriched as a consequence of the CDTX-induced muscle injury. Remarkably, “IL6 JAK STAT3 signaling” and “TNFA signaling via NF-kB” Hallmark gene sets were enriched in all possible pairwise comparisons including in the comparison SCI+CDTX vs all other 3 groups (Fig. 2, black arrows). The broader “Inflammatory response” gene set was enriched in all comparisons that included at least one group with CDTX muscular injury. Heat maps of these three gene sets also showed distinct expression patterns between SCI+CDTX and other three groups (Fig. 3). For initial validation, we confirmed that the *Osm* gene (encoding oncostatin M) in the “Inflammatory Response” gene set (which we have previously shown to be overexpressed in muscles following SCI+CDTX injury and playing a major role in SCI-NHO development^(20)^) was overexpressed in the SCI+CDTX group in the microarray analysis. We next selected candidate genes that were found upregulated in the SCI+CDTX group and known to play important roles in macrophage-mediated inflammatory responses such as *Csf1*, *Tnf*, *Il1b,* and *Ccl2.* Comparison 2 between SCI+CDTX group and Sham + CDTX group confirmed that “TNF / NF-κB signaling” and “IL6/JAK/STAT3” gene sets were significantly enriched in the SCI+CDTX group. The “inflammatory response” gene set also enriched (p=0.043) but the false discovery rate was 0.2 (Supplementary figure 1). Nevertheless, *Csf1*, *Tnf*, *Il1b, Ccl2*, and *Osm* mRNA were still among the leading-edge subsets (Supplementary figure 1C).

**Fig 3.**
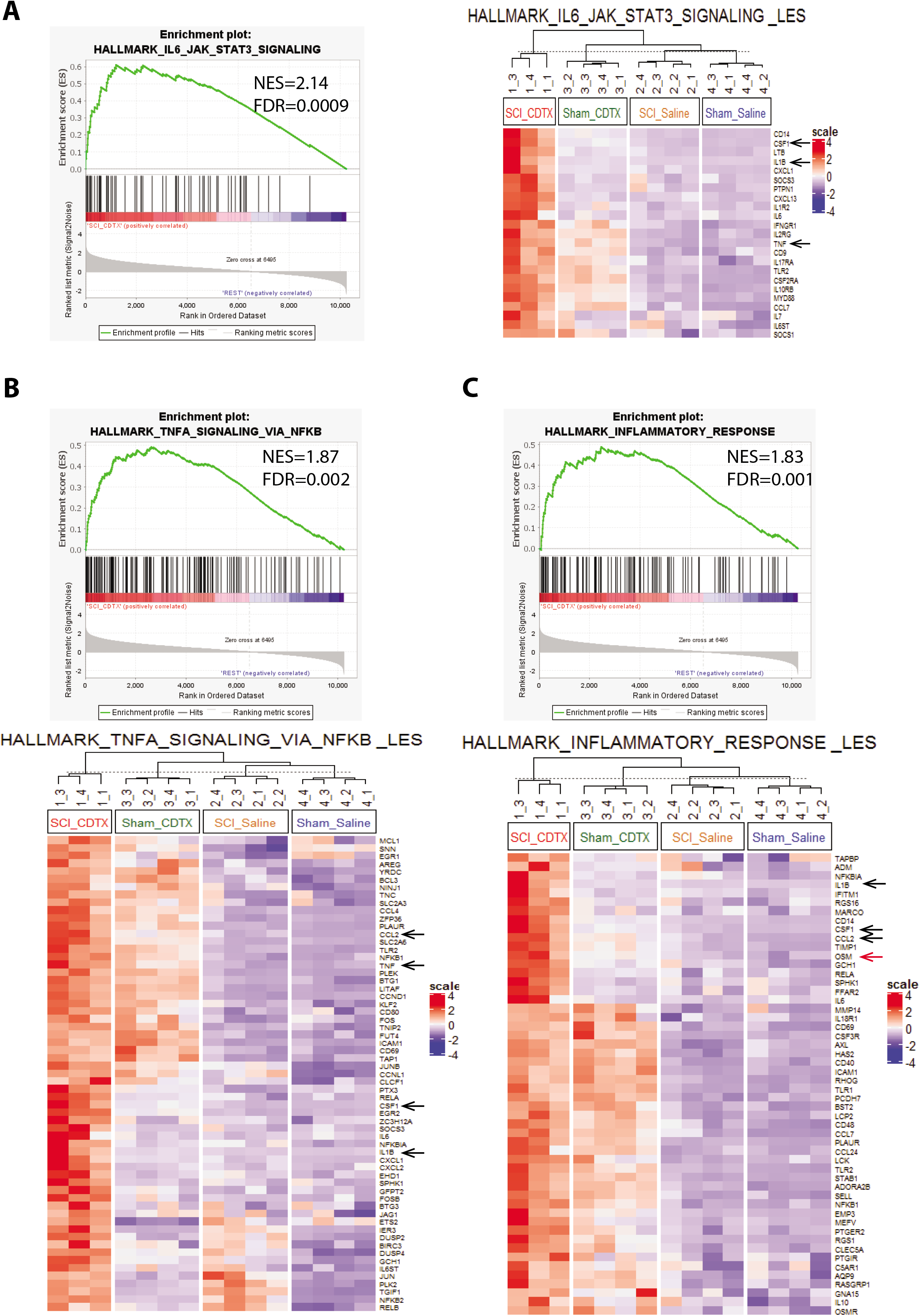
Heat maps of differentially expressed gene sets from injured muscles from mice that develop NHO following SCI and muscle injury. Enrichment score plots for (A) “IL6-JAK-STAT3 signaling”, (B) “TNF signaling via NFκB” and (C) “inflammatory response signaling” gene sets comparing injured muscles from SCI+CDTX group, which develops NHO, to the other 3 experimental groups that do not develop NHO. Heat maps of leading-edge subsets of for each of these gene sets is shown. Gene studied in subsequent experiments are indicated by black arrows. *Osm* previously studied in Torossian et al ^(20)^ is indicated by a red arrow.

Next, we validated these gene targets in another cohort of mice by quantitative real-time RT-PCR (qRT-PCR) (Fig. 4). CDTX injection increased the expression of *Csf1*, *Tnf*, *IL1b* and *Ccl2* two days post procedures regardless of SCI whereas *Il1b* expression was significantly higher in SCI+CDTX compared to Sham+CDTX similar to *Osm* ^(20)^. To further test the functional roles of CSF-1, TNF, CCL2 and IL-1β signaling pathways in SCI-NHO development *in vivo*, the four signaling pathways were blocked either by pharmacological inhibitors, neutralizing antibodies or using gene knockout mouse models.

**Fig 4.**
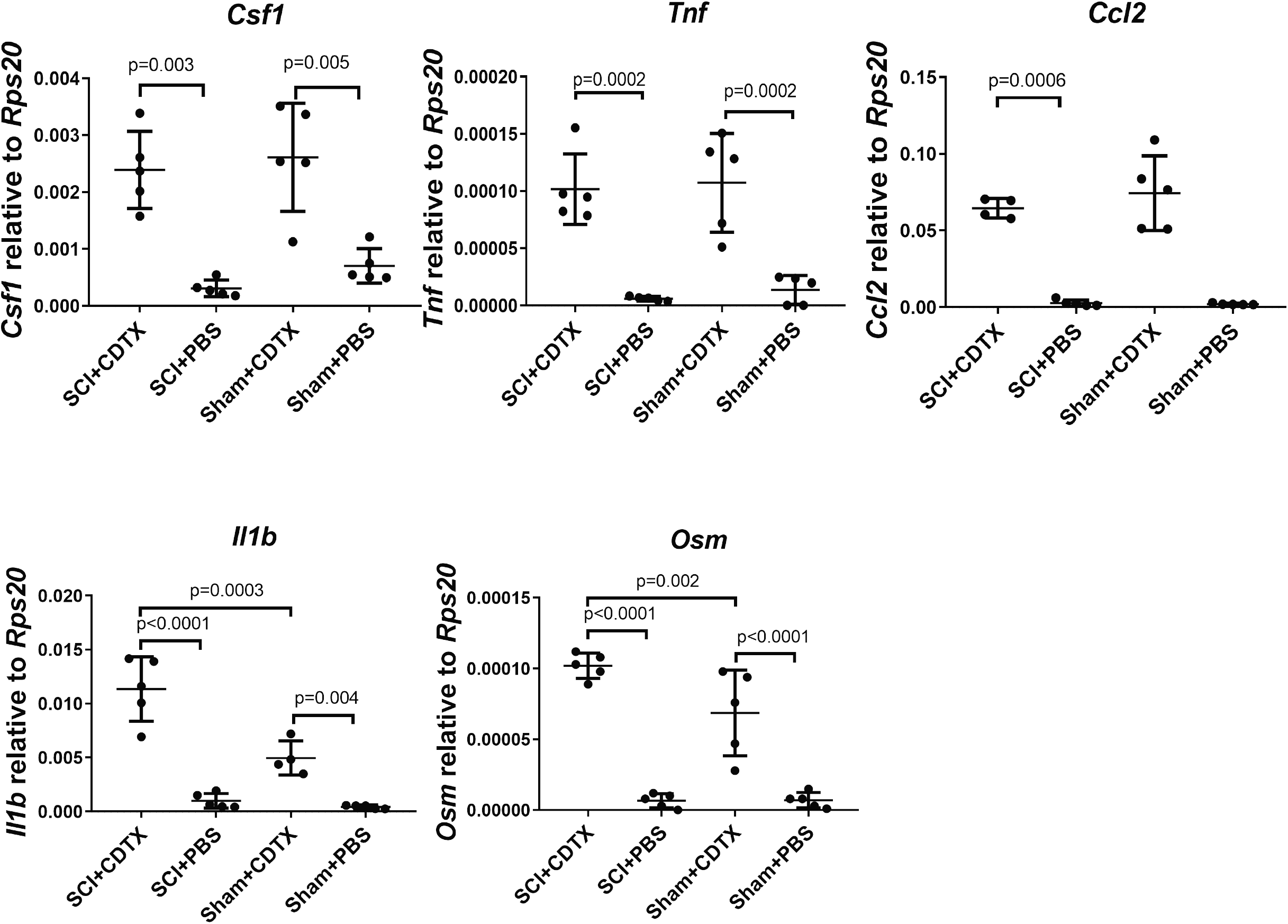
Validation of differentially expressed in injured muscles by qRT-PCR. *Csf1*, *Tnf*, *Ccl2*, *Osm* and *Il1b* mRNA expression in injured muscles from mice that underwent either SCI + CDTX muscle injury, SCI + PBS, Sham + CDTX or Sham + PBS. Muscles and total RNA were isolated two days post-surgery. Specified mRNAs were quantified relative to *Rps20* housekeeping gene. Each dot represents a separate mouse and bars are means ± SD (n=4-5/group). Statistical differences were calculated by ANOVA with Tukey’s multiple comparison test.

### CSF-1 receptor, TNF and CCR2 signaling do not promote NHO development

We first inhibited CSF-1 receptor signaling by administering GW2580, an orally bioavailable small inhibitor of CSF-1 receptor (CSF1R) tyrosine kinase ^(28,30)^. We first assessed the effect of GW2580 at inhibiting medullary monopoiesis stimulation induced by administration of a long-lived recombinant pig CSF-1-IgG Fc (pCSF1-Fc) fusion protein in vivo ^(29)^ (Supplementary Fig. 2). Flow cytometry analyses of BM cells showed that GW2580 abrogated increases in the numbers of CD11b^+^ F480^+^ Ly6G^-^ monocytes / macrophages in response to pCSF-1-Fc treatment and within this population, GW2580 was particularly effective at reducing the numbers of Ly6C^bright^ CSF1R^+^ and Ly6C^+^ CSF1Rlow monocytes whereas these treatments had no effect on the number of CD11b^+^ Ly6G^+^ granulocytes or CSF1R^-^ monocytes. Next, mice were administered daily GW2580 or vehicle from the day of SCI surgery and i.m. injection of CDTX (day 0) until day 10 and NHO volume were measured on day 10 by μCT *ex vivo*. Unexpectedly, GW2580 treatment significantly increased NHO volumes 1.6-fold following SCI (Fig. 5A). Thus CSF-1 is not a cytokine that promotes NHO development in the inflamed muscle following SCI.

**Fig 5.**
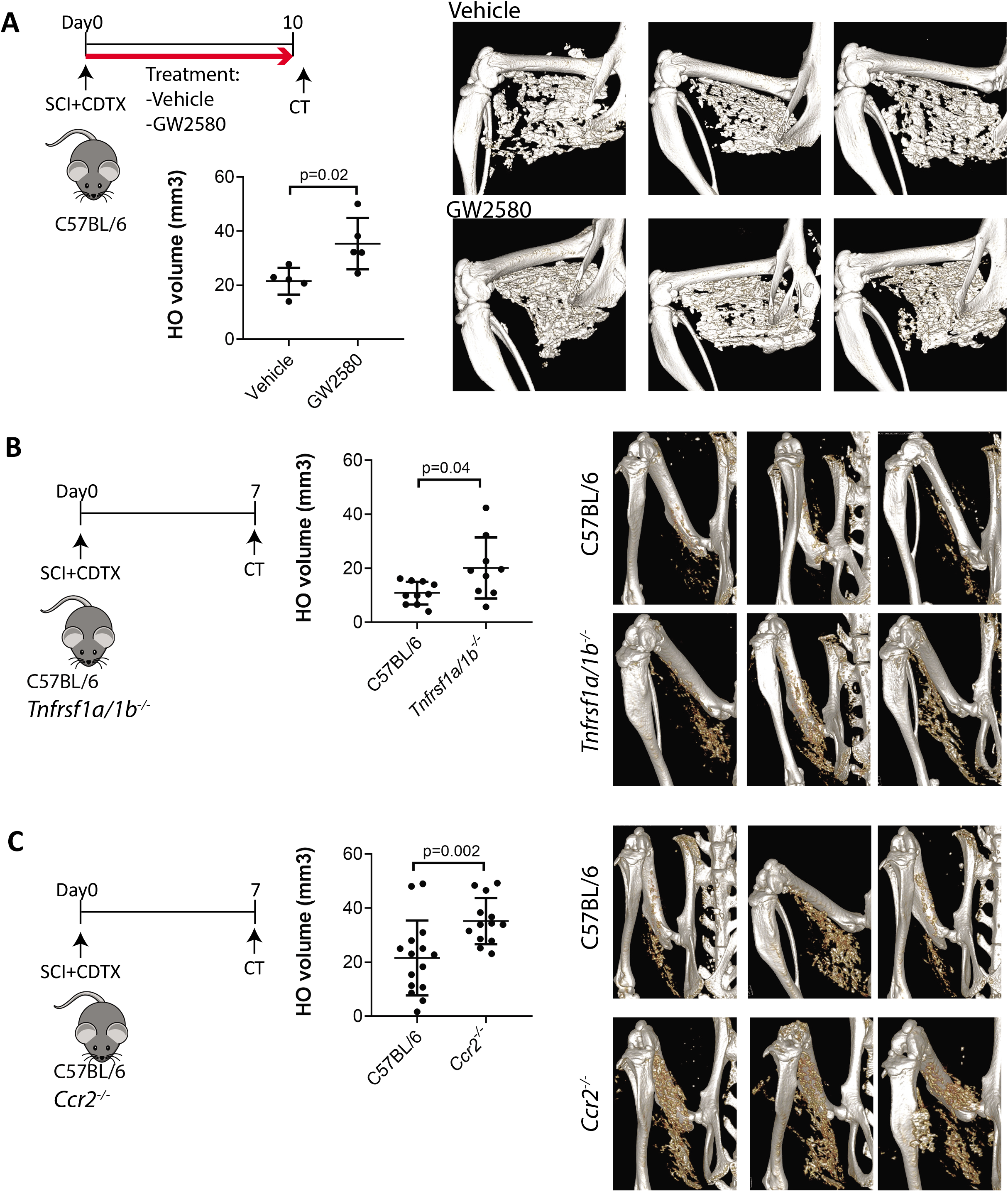
Blocking CSF1, TNF or CCR2 receptor signaling exacerbates NHO formation after SCI in mice. (A) C57BL/6 mice underwent SCI surgery and CDTX i.m. injection. Mice were then treated with vehicle or GW2580 (80mg/kg) twice daily from day 0 to 10. NHO formation was measured ex vivo by μCT on day 10 (n=5 mice/group). (B) *Tnfrsf1a/1b^−/−^* double knockout and C57BL/6 mice (n=10 / group from two separate experiments), (C) *Ccr2*^−/−^ and C57BL/6 mice (n= 14-15 mice / group from three separate experiments) underwent SCI surgery and CDTX i.m. injection. NHO formation was measured in μCT on day 7. Representative μCT 3D reconstitution images from each group and quantifications of NHO volumes for each mouse are shown. Each dot represents a separate mouse and bars are means ± SD. P values were calculated using two-sided Mann-Whitney test.

Next, to disrupt the TNF signaling pathway, we used mice with germinal deletion of the two genes encoding the two TNF receptors (*Tnfrsf1a/1b^−/−^* mice). Unexpectedly, NHO volume were 2-fold higher in in *Tnfrsf1a/1b^−/−^* double KO mice compared to C57BL/6 wild type mice (Fig. 5B). Therefore, similar to CSF-1, TNF signaling does not trigger NHO development after SCI but rather dampens it.

We have previously reported that infiltration of Ly6C^hi^ inflammatory monocyte/macrophages in the CDTX-injured muscles is significantly higher in mice with SCI compared to injured muscles from mice without SCI^(19)^. Furthermore, *in vivo* depletion of macrophages with clodronate-loaded liposomes reduced NHO formation dramatically ^(8)^. As CCL2/CCR2-mediated macrophage chemotaxis and recruitment largely contribute to inflammatory monocyte/macrophage infiltration in injured muscles ^(9,16)^, we further investigated whether defective CCR2-mediated signaling reduced NHO formation. Both CCR2-deficient (*Ccr2*^−/−^) and C57BL/6 wild-type mice underwent SCI surgery and i.m. CDTX injection and muscle leukocyte infiltrates were analyzed 4 days post-surgery. CCR2 deficiency led to a significant reduction in the infiltration of overall F4/80^+^ monocytes / macrophages as well as the Ly6C^hi^ subset. This was compensated in part by an increased recruitment of CD11b^+^ F4/80^-^ Ly6G^+^ neutrophils (Supplementary Fig. 3). Surprisingly however, NHO volumes were significantly increased 1.6-fold in *Ccr2*^−/−^ mice (Fig. 5C) despite reduced infiltration of inflammatory macrophages.

### IL-1-mediated signaling contributes to SCI-NHO development in mice

As IL-1 comprises two distinct but functionally redundant proteins, namely IL-1α and IL-1β encoded by two distinct genes, which both bind to their common IL-1 receptor IL1R1^(31,32)^, we tested the involvement of IL1-mediated signaling in *Il1r1*^−/−^ mice defective for IL1R1. *Il1r1*^−/−^ mice exhibited a significant 32.2% reduction in NHO volume compared to wild-type C57BL/6 control mice indicating IL-1 signaling play some roles in NHO development (Fig. 6A). However, when we conducted this experiment in mice defective for either IL-1α (*Il1a*^−/−^ mice) or IL-1β (*Il1b*^−/−^ mice), deficiency of neither IL-1β nor IL-1α alone affected NHO volumes (Supplementary Fig. 4). Interestingly, by qRT-PCR, we found that *Il1b* and *Il1a* mRNA expression changed over time. On day 2 post-injury, *Il1b* expression was significantly higher in SCI+CDTX compared to sham+CDTX groups, while it was significantly decreased from day 2 to day 4 (Figure 4, 6B). On the other hand, *Il1a* expression was significantly increased from day 2 to day 4 in SCI+CDTX muscles (Figure 6B). These results suggest different kinetics of induction of IL-1α and IL-1β in the injured muscle following SCI with potential functional redundancy. Therefore, we measured the effects of daily injections of combined neutralizing anti-mouse IL-1α and anti-mouse IL-1β monoclonal antibodies on NHO volumes. Antibody-mediated neutralization of both IL-1α and IL-1β caused a trend towards reduced NHO volumes although this did not reach statistical significance (Fig. 6C).

**Fig 6.**
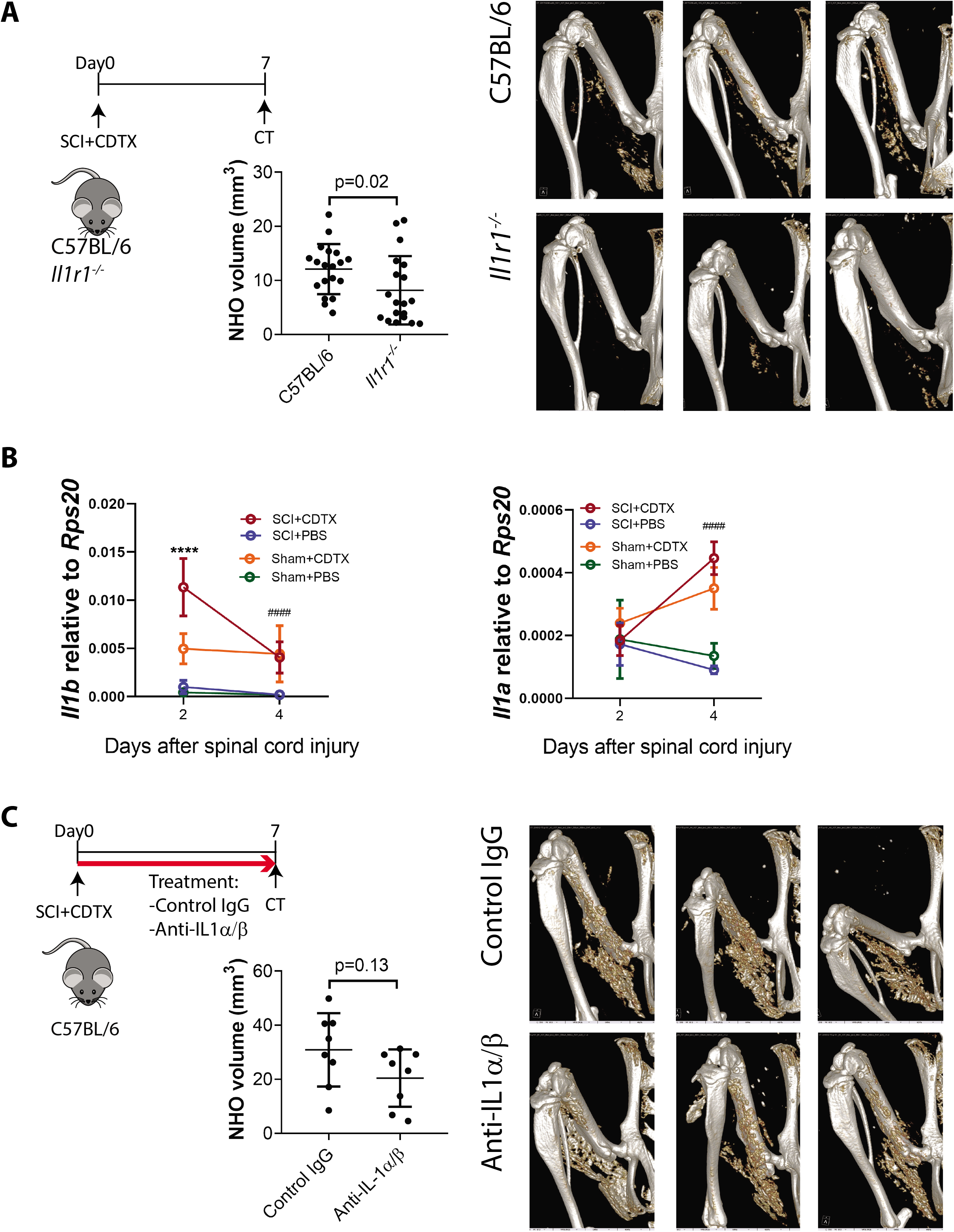
IL1 receptor deficiency reduces NHO formation after SCI in mice. (A) *Il1r1*^−/−^ and C57BL/6 mice underwent SCI surgery and i.m. injection of CDTX. NHO volumes were quantified by μCT 7 days post-injury. (B) qRT-PCR quantification of *Il1b* and *IL1a* mRNA in whole muscle from mice that underwent either SCI + CDTX, SCI + PBS, Sham + CDTX or Sham + PBS on day 2 and day 4 post-injury. The mRNA samples on day 2 are the same as figure 4. Expression of *Il1b* and *IL1a* were normalized to *Rps20* housekeeping gene. Data are presented as means ± SD (n=4-5/group). Statistical differences were calculated by two-way ANOVA with Sidak’s multiple comparison test. **** p<0.0001 SCI+CDTX vs Sham+CDTX on day2. #### p<0.0001 SCI+CDTX day2 vs day4. (C) C57BL/6 mice underwent SCI surgery plus i.m. injection of CDTX and injected daily with combination of anti-mouse IL1α and anti-mouse IL1β neutralizing Armenian hamster monoclonal antibodies or control Armenian hamster IgG. NHO volumes were quantified by μCT 7 days post-injury. Each dot represents a separate mouse and bars are means ± SD. P values were calculated using two-sided Mann-Whitney test. μCT scans of NHO formation are shown with representative images from each group.

### IL-1β protein is expressed in human NHO biopsies and increases NHO-derived FAP mineralization in vitro

To assess relevance to the human pathology, we studied IL-1β expression in human NHO biopsies taken from 13 patients by immunohistochemistry. Five patients had NHO following SCI and eight following TBI. There was a clear presence of CD68^+^ macrophages and multinucleated osteoclasts within NHO (Fig. 7A and Supplementary Fig. 5) as we previously described^(8,20)^. We noted IL-1β^+^ CD68^+^ mononucleated macrophages adjacent to the compact bone of NHOs, and also distributed amongst the inflamed fibrotic tissue surrounding NHOs (Fig. 7A, Supplementary Fig. 5). Interestingly, for four biopsies we observed the presence of IL-1β^+^ CD68^+^ multinucleated osteoclasts (Fig. 7A, Supplementary Fig. 5). We also confirmed that patients with NHOs had significantly elevated plasma IL-1β concentration compared to healthy donors (Fig. 7B) suggesting that IL1-β may play a role in NHO development.

**Fig 7.**
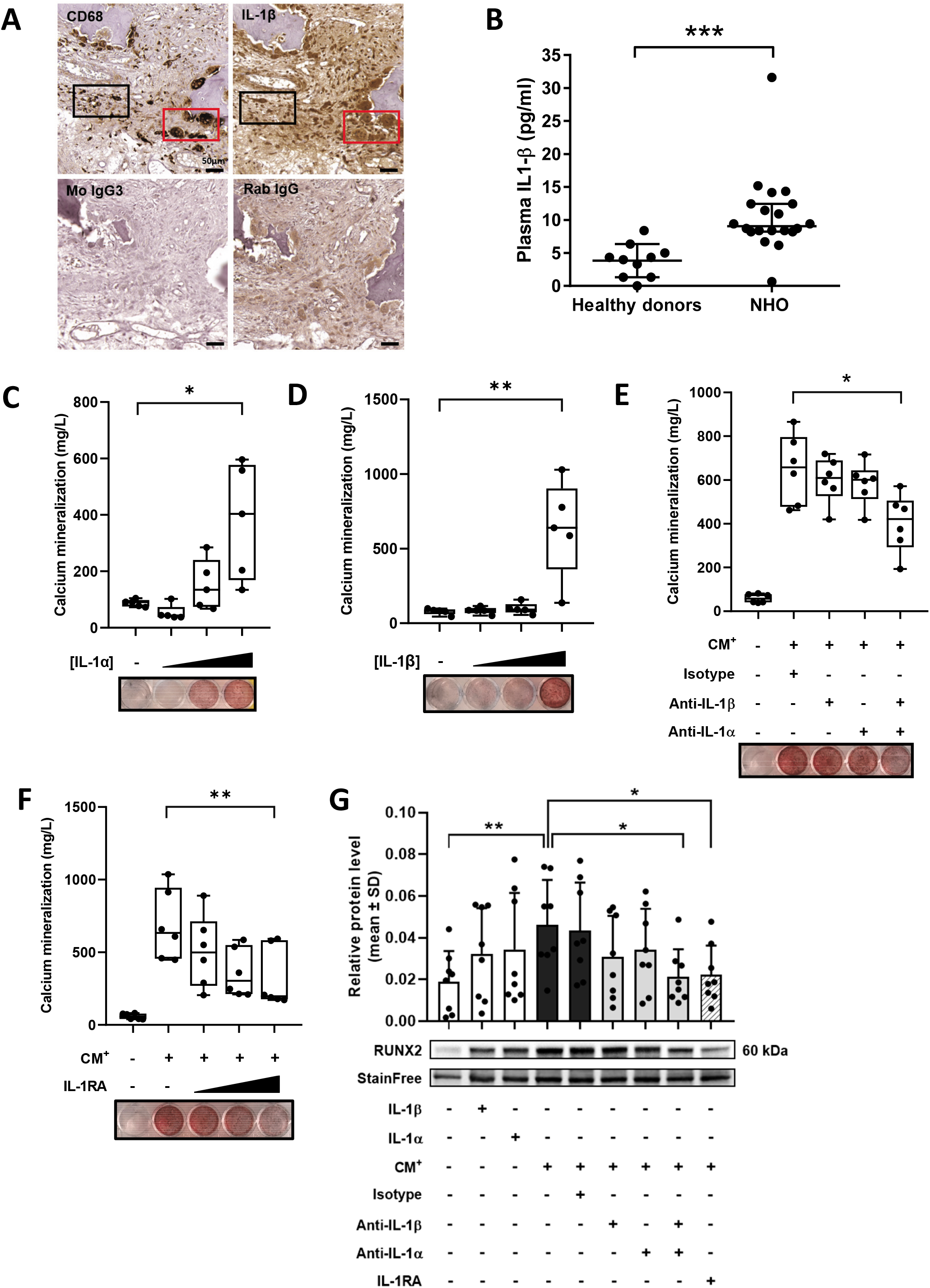
IL-1β is expressed in human NHO biopsies and stimulates calcium mineralization of NHO-derived FAPs *in vitro*. (A) Representative immunohistochemistry staining of serial sections of a human NHO biopsy (SCI patient) demonstrating the colocalization of the macrophage marker CD68 (top panel left) and IL-1β protein expression (top panel right). Black rectangle shows IL-1β expressing CD68^+^ macrophages in the fibrotic tissue surrounding NHO while red rectangle shows IL-1β expressing multinucleated CD68^+^ osteoclasts. Mouse IgG3 (bottom left panel) and Rabbit IgG (bottom right panel) confirm the specificity of CD68 and IL-1β antibodies. 20X magnification and scale bar = 50μm. (B) IL-1β concentration in plasma of healthy volunteers and NHO patients. Each dot represents a different individual. Bars are means ± SD. Statistical difference was calculated using two-sided non-parametric Mann-Whitney test, *** p<0.001. (C) Dose-response of recombinant human IL-1α (0.01; 1 and 100 ng/mL) and (D) IL-1β (0.01; 0.5 and 10 ng/mL) on calcium mineralization of PDGFRα^+^ CD56^-^ FAPs derived from 5 different NHO biopsies / patients cultured in osteogenic medium. (E) Osteogenic differentiation assay of FAPs cultured with LPS-stimulated CD14^+^ blood monocyte-conditioned medium (CM^+^) and incubated with anti-IL-1α, anti-IL-1β neutralizing antibodies or isotype control (100 ng/mL). (F) Osteogenic differentiation assay of FAPs derived from 6 different biopsies / patients cultured with CM^+^ and incubated with 10, 100 or 500 ng/mL of recombinant human IL1RA. In (C-E), each dot represents a result from a different NHO biopsy / patient. Boxes extend from the 25th to 75th percentiles, middle line and whiskers represent respectively median and minimum and maximum value. Statistical differences were analyzed using non-parametric repeated measure Friedman test with Dunn’s correction for multiple comparisons, * p < 0.05; ** p < 0.01. (G) RUNX2 protein expression level of FAPs evaluated by Western blot following 7 days of osteogenic differentiation with IL-1β (10 ng/mL), IL-1α (100 ng/mL), NHOmac CM^+^, control isotype antibody, anti-IL-1α and anti-IL-1β antibody (100 ng/mL) or IL-1RA (500 ng/mL). Statistical differences were analyzed using non-parametric repeated measure Friedman test with Dunn’s correction for multiple comparisons, * p < 0.05; ** p < 0.01.

To further investigate the involvement IL-1 on NHO pathology, we collected surgical residues of muscle tissue surrounding NHO after resection surgery. Muscle progenitor cells were expanded *in vitro* and PDGFRα^+^ CD56^-^ FAPs were isolated by FACS (Supplementary Fig. 6). Osteogenic differentiation assays showed that FAPs exhibited higher calcium mineralization activity when cultured with increasing doses of either recombinant IL-1α (Fig. 7C) or IL-1β (Fig. 7D) suggesting that IL-1 stimulates FAP osteoblastic differentiation. The addition of anti-IL-1β neutralizing antibody significantly inhibited FAP calcium mineralization activity when cultured with IL-1β (Supplementary Fig. 7A). Considering the pivotal role of macrophages in NHO formation, we isolated CD14^+^ blood monocytes from healthy donors and NHO-bone marrow derived CD14^+^monocytes. CD14^+^ cells were cultured with or without LPS to generate CM^-^ and LPS-stimulated CM^+^ conditioned media. CM^+^ displayed significantly higher IL-1α and IL-β concentrations compared to CM^-^ (Supplementary Fig. 7B and 7C) and, as expected, pooled CM^+^ stimulated FAPs calcium mineralization activity in osteogenic differentiation assays (Fig. 7E and 7F). When added individually, anti-IL-1α and anti-IL-1β neutralizing antibodies did not significantly reduce calcium mineralization of CM^+^-stimulated FAPs, however, the combination of both antibodies significantly reduced FAPs mineralization activity associated with significant decrease in RUNX2 protein expression (Fig. 7E and 7G), suggesting a synergistic effect of IL-1α and IL-1β. This was further confirmed by the dose-dependent inhibitory effect of IL-1RA, the endogenous protein antagonist of IL1R1 common receptor for IL-1α and IL-1β, on FAPs mineralization associated with reduced RUNX2 protein expression (Fig. 7F and 7G). Overall, these results confirm the involvement of IL-1 as inflammatory mediator of NHO formation in the human pathology.

## Discussion

Using non-supervised unbiased GSEA of whole muscle RNA in mice, we demonstrate herein that superimposition of a SCI to a muscle injury exacerbates the local inflammatory response to muscle injury, and most particularly expression of genes associated with IL6 cytokine family / JAK / STAT3 signaling and TNF signaling via NF-κB. The proinflammatory effect of the SCI is rapid as we measured gene expression 2 days after SCI and /or muscle injury. This is consistent with our previous study showing that SCI causes persistent and abnormally high expression of the IL6 family cytokine OSM, together with persistently high levels of active tyrosine phosphorylated STAT3 via JAK1/2 tyrosine kinase activation in the injured muscle ^(19)^. In addition to OSM that we have already shown to promote NHO in response to SCI ^(20)^, we also find that SCI increased the expression of other inflammatory cytokines in injured muscles including TNF, CSF-1 and IL-1β as well as inflammatory chemokines such as CCL2, CXCL1 or CXCL13 (Fig. 3). As we have previously shown that macrophages play a key role in NHO development ^(8)^, we focused on TNF, CSF-1 and CCL2 which are well known to regulate macrophage polarization and recruitment ^(33–36)^ as well as IL-1β, which is specifically produced by phagocytes following activation of inflammasomes and mediates inflammatory responses in many tissues ^(37)^. Using either mice lacking the cognate cytokine receptors for TNF, CCL2 and IL-1, neutralizing antibodies (IL-1), or small receptor antagonists (CSF-1), we found that of these 4 potential mediators, only IL-1 promoted NHO development after SCI. Germinal deletion of the *Il1r1* gene that encodes the common receptor for IL-1α and IL-1β significantly reduced NHO volumes following SCI although this decrease was not as pronounced as observed in *Osmr*^−/−^ mice lacking the OSM receptor ^(20)^. Active IL-1β release by macrophages is a 2-step process, first requiring activation of NF-κB-dependent transcription of the *Il1b* gene followed by inflammasome-dependent activation of non-active IL-1β pro-peptide by caspase-1-mediated proteolytic cleavage. In contrast the IL-1α pro-peptide is biologically active and pre-formed intracellularly in many cell types, particularly endothelial cells and fibroblasts and its activity can be further enhanced by calpain and several caspases upon cell death ^(38)^. Therefore while IL-1β needs two distinct signals to be produced in active form by macrophages, pre-formed IL-1α is a “pre-loaded gun” that is released upon death of many cell types ^(37,38)^. Although we could not detect enhanced IL-1α transcription in our gene expression microarray analysis at day 2 post-injury, *Il1a* was upregulated later at day 4 and it is likely that muscle destruction caused by CDTX injection would release IL-1α. To assess the respective role of IL-1α and IL-1β in NHO pathogenesis, we used mice targeted deletion of either the *Il1a* gene or *Il1b* gene. Deletion of either IL-1 gene alone did not alter NHO development suggesting that both IL-1 proteins act in concert to promote NHO development. This is further suggested by reduced NHO development in mice treated simultaneously with two neutralizing antibodies specific for IL-1α and IL-1β although this decrease did not reach significance (p=0.13, Fig. 6B).

To validate our results in the human pathology, we show that IL-1β protein is present in the NHO itself and the non-ossified and peripheral part of the particularly by CD68^+^ macrophages. It indicates that IL-1β is produced locally by activated macrophages and could contribute to the development of NHO. Furthermore, IL-1β protein concentration was doubled in the plasma of SCI patients developing NHO compared to healthy subjects. Polynucleated osteoclasts were also identified for several NHO biopsies suggesting active bone remodeling. Functionally, we find that purified recombinant IL-1α and IL-1β both potently promote calcium deposition and mineralization of human FAPs purified from the muscles surrounding patient NHOs. Furthermore, we have had previously shown that LPS-stimulated monocyte-conditioned medium (CM^+^) promotes human FAP osteogenic differentiation and calcium mineralization in part via OSM secretion ^(20)^. Herein we show that IL-1α and IL-1β, which are both secreted by human monocytes in response to LPS, both contribute to the pro-osteogenic effect of CM^+^ on human FAPs. Similar to our results suggesting that antibody neutralization of both IL-1α and IL-1β is necessary to reduce NHO development in mice *in vivo*, neutralization of both IL-1α and IL-1β was necessary to reduce the pro-osteogenic effect of CM^+^ on human FAPs *in vitro*, an effect recapitulated by addition of recombinant IL1RA, the physiological endogenous antagonist of the IL-1 receptor. As we have shown that muscle FAPs are the cells-of-origin of NHO in both humans and mice (Tseng HW et al, revised manuscript submitted), this suggests that both IL-1α and IL-1β are potential mediators of NHO development in patients.

In respect to the other inflammatory cytokines and chemokines that we found to be highly expressed in muscles developing NHOs in mice, namely TNF, CCL2 and CSF-1, we unexpectedly found that they did not promote but instead dampened NHO development following SCI. In respect to TNF and CCL2, both have been shown to play a major role in normal muscle repair by promoting apoptosis of FAPs proliferating in the first few days in injured skeletal muscles and are therefore essential to prevent muscle fibrosis ^(16)^. As FAPs are the cells-of-origin of NHO in both humans and mice (Tseng HW et al, revision submitted), we speculate that *Ccr2*^−/−^ and *Tnfrsf1a/1b^−/−^* mice may have reduced apoptosis and prolonged proliferation of FAPs ^(16)^, which could be responsible for increased muscle fibrosis and NHO volumes. The enhancing effect of CSF1R tyrosine kinase inhibitor GW2580 on NHO development was also surprising because CSF1R signaling is necessary to maintain many tissue-resident macrophages and osteoclasts but dispensable to monocyte production in the bone marrow ^(39,40)^ although extra-physiological treatment with stable CSF-1 protein considerably enhances monopoiesis ^(29,40)^. As macrophages are necessary to the development of SCI-induced NHO ^(8)^ as well as genetically-driven HO development in *fibrodysplasia ossificans progressiva* ^(17,41)^, we anticipated that inhibition of CSF1R signaling would reduce NHO formation. The opposite effect that we observed on NHO formation *in vivo* may be due to the complex function of CSF-1 on macrophage biology. CSF-1 is not only necessary to the maintenance of many types of tissue-resident macrophages, it also induces and modulates macrophage pro-inflammatory or anti-inflammatory polarization in a context-dependent manner ^(35,42)^. Our results herein suggest that CSF-1 polarizes macrophages in an inflammatory manner in injured muscles in the context of a SCI similar to the effect of GW2580 treatment or CSF-1-defiency which both inhibit functional recovery and increase tubulointerstitial fibrosis in a mouse model of acute kidney injury ^(43,44)^.

Although our mouse model of SCI-NHO recapitulates many features of NHO observed in patients with SCI ^(8)^, a limitation of our mouse model is that it does not recapitulate the endochondral phase observed within NHO from patients. Indeed, safranin-O staining showed a clear chondrogenic phase with different stages of cartilage and chondroblast maturation in NHO biopsies resected from patients with SCI suggesting some endochondral bone formation. However, we could not detect any chondrogenic phase in the NHO nodules forming in the injured muscles in mice that underwent SCI (Supplementary Fig. 8) suggesting intramembranous bone formation. Despite this limitation, functional in vivo experiments in mice, immunohistology and functional in vitro experiments with biopsies from patients suggest a commonality of triggering mechanisms involving exacerbated inflammation in muscles developing NHO ^(8)^ with a role of OSM ^(20)^ and IL-1 reported herein.

In conclusion, we report herein that SCI exacerbates the transcription of several inflammatory cytokines and chemokines such as IL-1β, IL-1α, TNF, CSF-1 and CCL2 in injured muscles. We established that TNF, CSF-1 and CCL2 had a dampening effect on NHO development. In sharp contrast, IL-1 signaling promoted NHO development in injured muscles in our mouse model of SCI-induced NHO. The expression of IL-1β in NHO biopsies from SCI and TBI patients and the osteogenic effect of IL-1α and IL-1β secreted by activated monocytes was confirmed in cultures of FAPs isolated from the muscles surrounding NHO biopsies from SCI and TBI patients. These results are consistent with the observation that 80% children with the extremely rare genetic deficiency of IL-1 receptor antagonist (DIRA) syndrome, in which IL-1 activity cannot be inhibited by endogenous IL1RA, develop periarticular HO ^(45)^. Considering that both OSM ^(20)^ and IL-1 (herein) promote NHO development following SCI, prophylactic combinations of drugs antagonizing IL-1 signaling (such as anakinra or IL1RA) and JAK1/2 tyrosine kinases (such as ruxolitinib) activated by the OSM receptor^(19)^ may be studied in more detail to assess their potential synergism in inhibiting NHO development. How SCI causes this peculiar exacerbation of inflammation in injured muscles that leads to NHO formation remains incompletely understood. In our SCI-NHO mouse model, transection of the spinal cord between T11-T13 leads to a significant increase of epinephrine blood plasma concentration 24 hours post-surgery and we have shown chemical sympathectomy or inhibition of β1/2-adrenergic receptors with propranolol significantly reduced NHO formation, suggesting that sympathetic adrenergic signaling plays an important role in NHO pathogenesis ^(46)^. By the same token, the increased adrenergic signaling subsequent to a SCI may exacerbate the inflammatory response in injured muscle. In support of this, binding of epinephrine to macrophages via β2 adrenergic receptors has been shown to modulate inflammatory responses by for instance up-regulating OSM and IL-1β expression in macrophages and microglial cells ^(47–49)^. A similar mechanism may occur via systemic release of epinephrine from adrenal glands in response to low level SCI ^(46)^.

## Acknowledgements

This research was funded in parts by Project Grant 1101620 (JPL and FG) and Ideas Grant 1181053 (JPL, KAA, HWT, FG, SB) from the National Health and Medical Research Council of Australia (NHMRC), the Congressionally Approved Spinal Cord Injury Research Program of the US Department of Defense award W81XWH-15-1-0606 (JPL and FG), The University of Queensland Early Career Researcher Grant UQECR1834805 (KA) and by funds from the Mater Foundation. JPL and KS supported by Research Fellowships 1044091 and 1141131 from the NHMRC. IK was supported by an international student scholarship from The University of Queensland. MS was supported by a student scholarship from the French Society of Physical and Rehabilitation Medicine.

The authors also greatly acknowledge the help and advice of Dr Adam Ewing and Dr Kim Summers (Mater Research Institute) and the technical assistance of Marie-Emmanuelle Goriot, Bastien Rival (IRBA, INSERM UMRS-MD-1197, Clamart), Denis Clay (INSERM UMS-44 Université de Paris-Saclay, Villejuif) and Guillaume Genêt (Raymond Poincaré Hospital, Garches). The authors also acknowledge the technical assistance of “Cochin HistIM Facility” (Paris). The authors also acknowledge the scientific and technical assistance at the Translational Research Institute: The University of Queensland biological resources TRI facility, histology facility, flow cytometry facility, microscopy facility and the preclinical imaging facility, which is supported by Therapeutic Innovation Australia (TIA). TIA is supported by the Australian Government through the National Collaborative Research Infrastructure Strategy (NCRIS) program.

Authors’ contributions: HWT, KAA, IK, DG, JG, CV, MS, WF, BJ, SMM, GT, LW and CV performed the experiments. HWT, KAA, IK, DG, SB and JPL conceived the experiments and the work. KS provide essential reagents and expertise in IL1 mouse models. HWT, KAA and JPL wrote the manuscript. HWT, KAA, DG, FG, KS, SB and JPL edited the manuscript. HWT, KAA, FG, SB and JPL obtained funding.

