## Supplementary materials for "Interleukin-1 is overexpressed in injured muscles following spinal cord injury and promotes neurogenic heterotopic ossification"

A

B

C

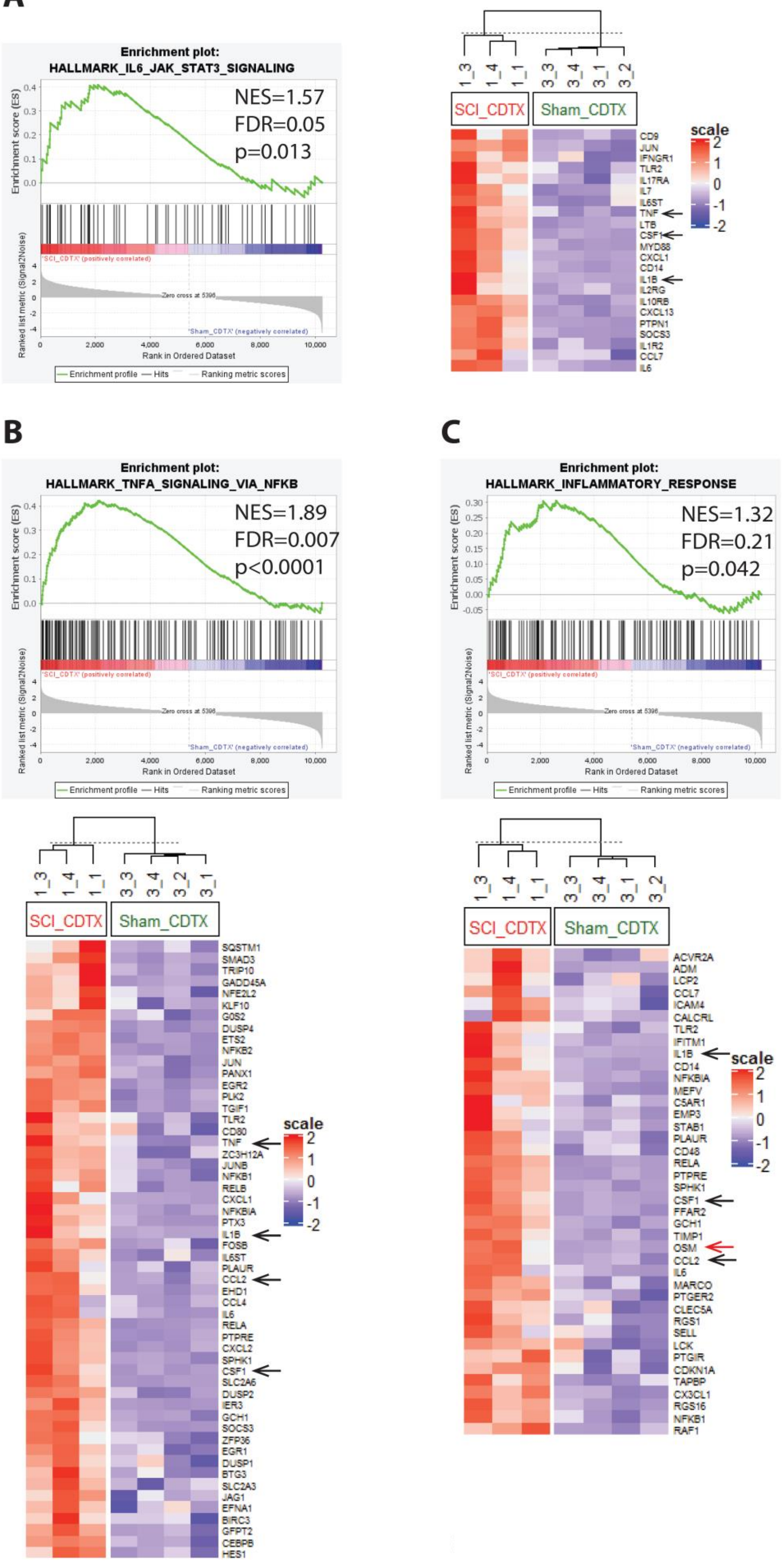
Enrichment plot:
HALLMARK_IL6_JAK_STAT3_SIGNALING
NES=1.57
FDR=0.05
p=0.013
Enrichment score (ES)
Ranked list metric (Signal2Noise)
'SCI_CDTX' (positively correlated)
Zero cross at 5396
'Sham_CDTX' (negatively correlated)
Rank in Ordered Dataset
Enrichment profile
Hits
Ranking metric scores
1_3
1_4
1_1
3_3
3_4
3_1
3_2
SCI_CDTX
Sham_CDTX
scale
CD9
JUN
IFNGR1
TLR2
IL17RA
IL7
IL6ST
TNF
LTB
CSF1
MYD88
CXCL1
CD14
IL1B
IL2RG
IL10RB
CXCL13
PTPN1
SOCS3
IL1R2
CCL7
IL6
Enrichment plot:
HALLMARK_TNFA_SIGNALING_VIA_NFKB
NES=1.89
FDR=0.007
p<0.0001
SQSTM1
SMAD3
TRIP10
GADD45A
NFE2L2
KLF10
G0S2
DUSP4
ETS2
NFKB2
JUN
PANX1
EGR2
PLK2
TGIF1
TLR2
CD80
TNF
ZC3H12A
JUNB
NFKB1
RELB
CXCL1
NFKBIA
PTX3
IL1B
FOSB
IL6ST
PLAUR
CCL2
EHD1
CCL4
IL6
RELA
PTPRE
CXCL2
SPHK1
CSF1
SLC2A6
DUSP2
IER3
GCH1
SOCS3
ZFP36
EGR1
DUSP1
BTG3
SLC2A3
JAG1
EFNA1
BIRC3
GFPT2
CEBPB
HES1
Enrichment plot:
HALLMARK_INFLAMMATORY_RESPONSE
NES=1.32
FDR=0.21
p=0.042
ACVR2A
ADM
LCP2
CCL7
ICAM4
CALCRL
TLR2
IFITM1
IL1B
CD14
NFKBIA
MEFV
C5AR1
EMP3
STAB1
PLAUR
CD48
RELA
PTPRE
SPHK1
CSF1
FFAR2
GCH1
TIMP1
OSM
CCL2
IL6
MARCO
PTGER2
CLEC5A
RGS1
SELL
LCK
PTGIR
CDKN1A
TAPBP
CX3CL1
RGS16
NFKB1
RAF1

**Supplementary Fig. 1.** Gene sets enriched in the injured muscles developing NHO in mice with SCI (SCI_CDTX) compared to injured muscles in mice without SCI (Sham_CDTX). Enrichment score plots and heat maps of leading-edge subsets for (A) “IL6-JAK-STAT3 signaling”, (B) “TNF signaling via NFκB” and (C) “inflammatory response signaling” gene sets comparing injured muscles from SCI+CDTX group to Sham+CDTX group that do not develop NHO. Genes investigated in the present study are indicated by black arrow. *Osm* previously studied in Torossian et al [20] is indicated by a red arrow.

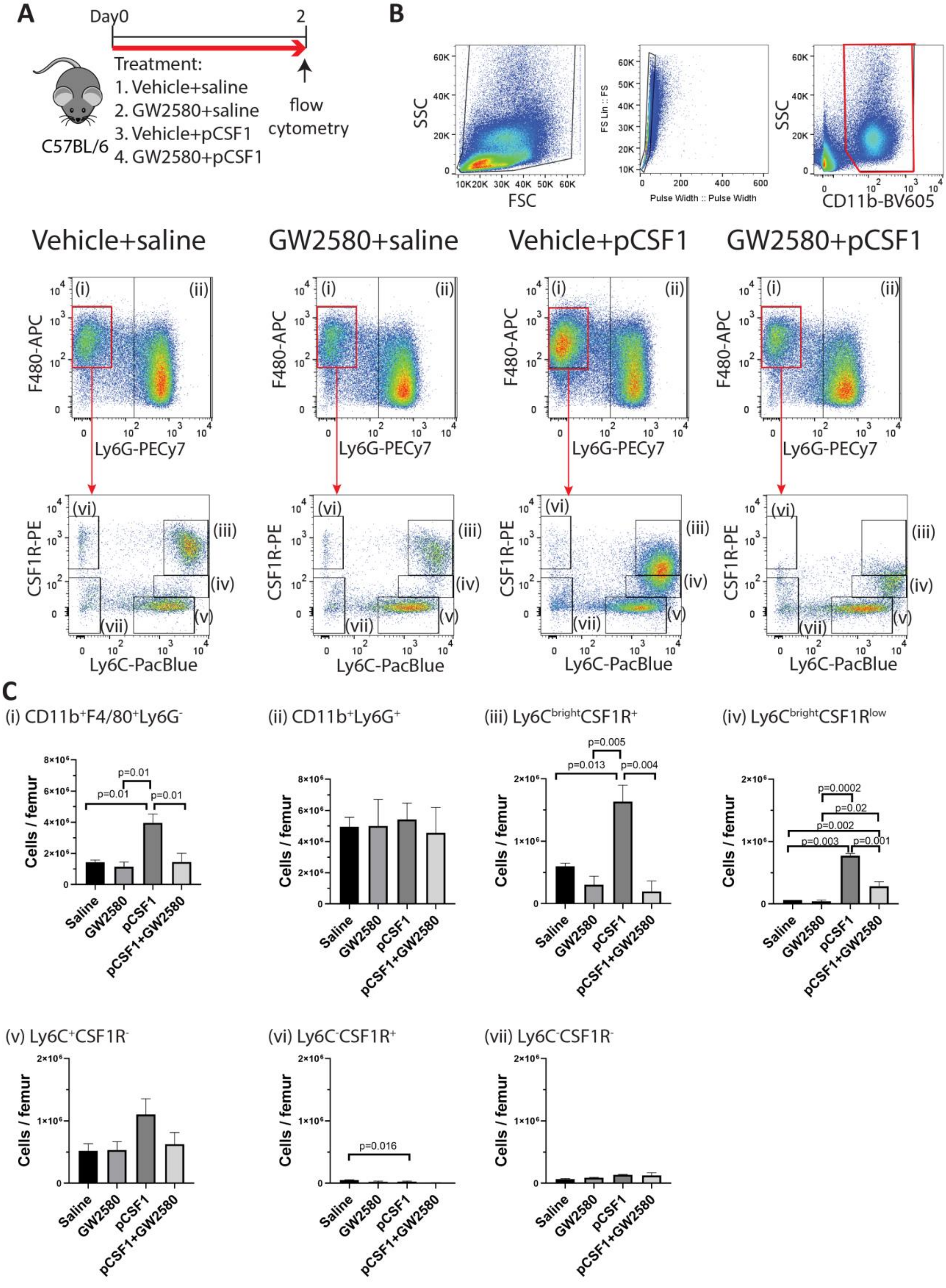


Supplementary Figure 2

**Supplementary Fig 2.** In vivo effects of GW2580 treatment on CSF-1-induced monopoiesis in mice. (A) Schematic of the experimental groups. C57BL/6 mice were injected with pigCSF1-IgGFc (pCSF1) or saline and gavaged with GW2580 or vehicle for 2 consecutive days and BM harvested on the following morning. (B) Gating strategy to identify BM CD11b$^+$ Ly6G$^+$ granulocytes (ii), CD11b$^+$ F4/80$^+$ Ly6G$^-$ monocytes / macrophages and their subsets according to Ly6C and CSF1R expression (iii-vii). (C) Absolute numbers of granulocytes and monocyte / macrophage subsets in the femoral BM of mice treated with vehicle, pCSF1 and/or GW2580. Data are means ± SD of 2 mice per group. P values were calculated by ANOVA with Tukey's multiple comparison test.

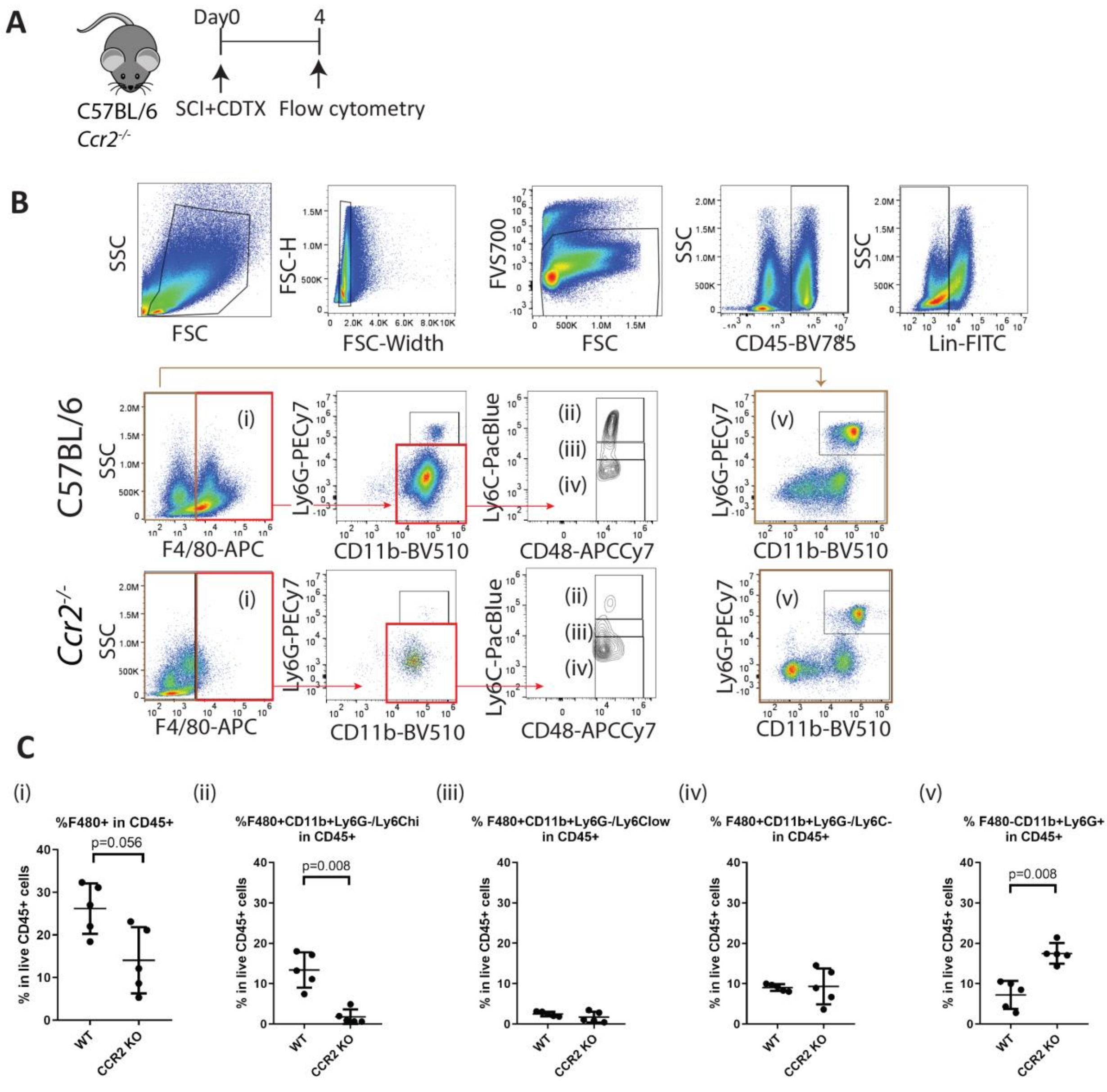


Supplementary Figure 3

**Supplementary Fig 3.** CCR2 deficiency reduces inflammatory monocyte infiltration in injured muscles after SCI. (A) C57BL/6 control and *Ccr2*$^{-/-}$ mice underwent SCI and i.m. injection of CDTX (n=5 mice per group). Leukocytes were isolated from injured muscles 4 days post injury and cell populations were analyzed by flow cytometry. (B) After gating out FVS700$^{+}$ dead cells, CD45$^{+}$ and Lineage (CD45R/B220, Ter119, CD3ε)-negative leukocytes were gated as F4/80$^{-}$ CD11b$^{+}$ Ly6G$^{+}$ neutrophils (v) and F4/80+ CD11b+ Ly6G-monocyte / macrophages (i) that were further subgated according to CD48 and Ly6C expression (ii-iv). Cell surface expression of CCR2 was also examined on monocyte macrophage subsets ii-iv. (C) Frequencies of granulocytes and monocyte / macrophage subsets in the injured muscles. Each dot represents a separate mouse and bars are mean ± SD. P values were calculated with two-sided Mann-Whitney test.

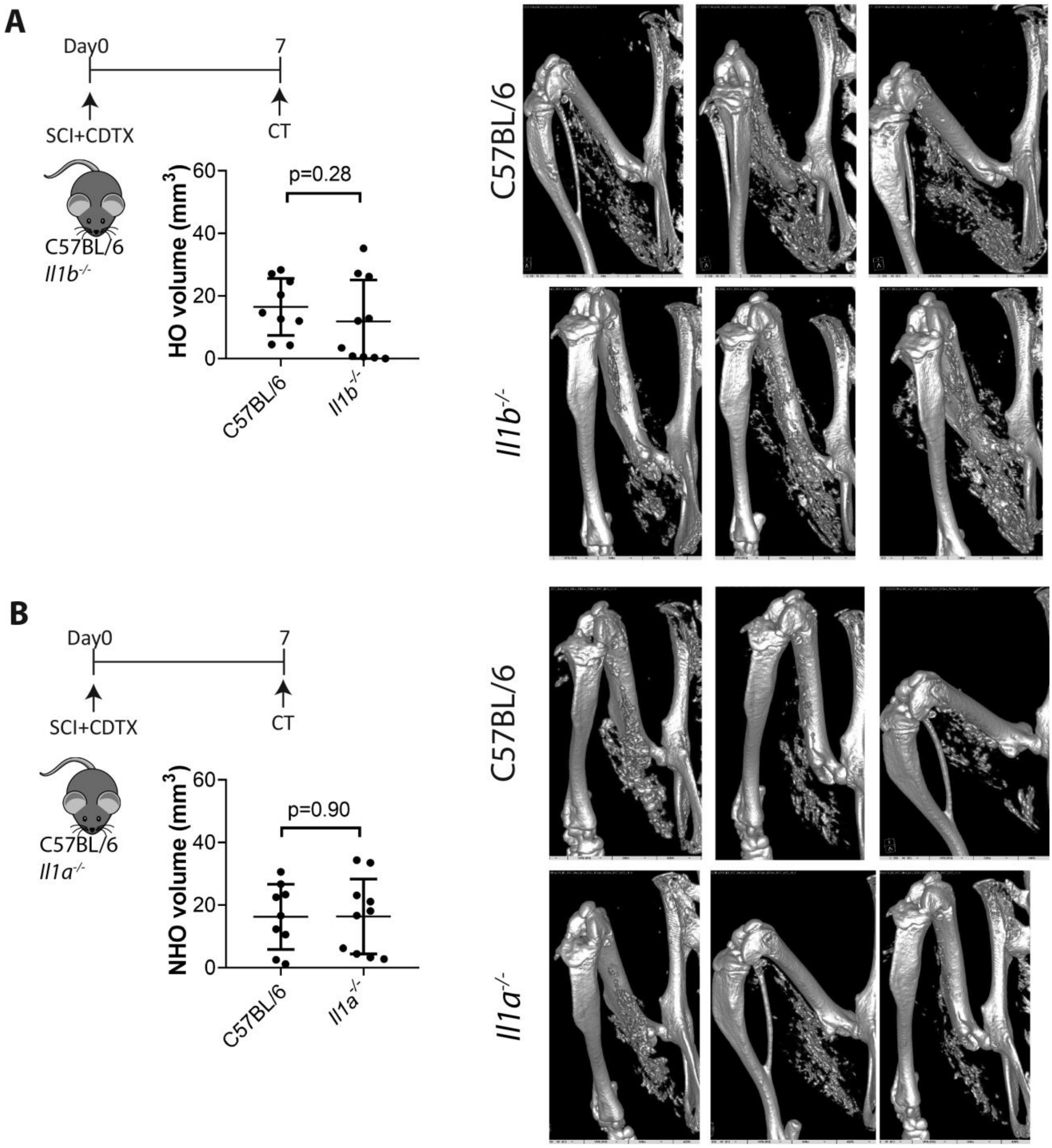


**Supplementary Fig 4.** Deficiency in either IL-1α or IL-1β does not impact NHO formation. (A) *Il1a*$^{-/-}$ mice or (B) *Il1b*$^{-/-}$ mice and wild-type control C57BL/6 mice underwent SCI surgery and i.m. injection of CDTX. NHO volume was quantified on µCT 7 days post-injury. NHO volumes were quantified by µCT 7 days post-injury. Data are means ± SD with each dot representing an individual mouse. P values were calculated using two-sided Mann-Whitney test. NHO formation are shown with representative images from each group.

**A**

| | $CD68^+$macrophages | $CD68^+IL\text{-}1\beta^+$ macrophages | Presence of $CD68^+IL\text{-}1\beta^+$ osteoclasts |
|---|---|---|---|
| SCI 1 | ++ | ++ | Y |
| SCI 2 | +/- | +/- | N |
| SCI 3 | ++ | + | N |
| SCI 4 | +++ | ++ | N |
| SCI 5 | ++ | ++ | Y |
| TBI 1 | + | + | N |
| TBI 2 | +/- | +/- | N |
| TBI 3 | ++ | ++ | Y |
| TBI 4 | +/- | +/- | N |
| TBI 5 | ++ | + | N |
| TBI 6 | ++ | ++ | N |
| TBI 7 | ++ | - | N |
| TBI 8 | ++ | ++ | Y |

**B**

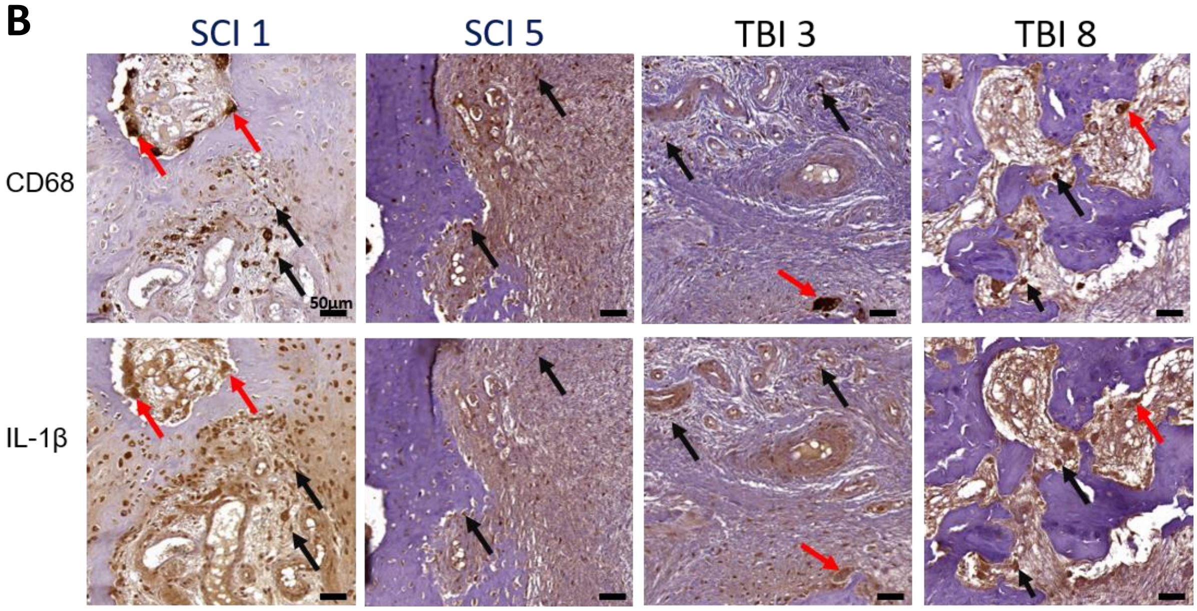


**Supplementary Fig 5.** Overview of CD68 and IL-1β immunohistochemistry staining of human NHO biopsies. (A) Table recapitulating the presence of $CD68^+$ expressing macrophages, $CD68^+$ IL-$1\beta^+$ expressing macrophages and $CD68^+$ IL-$1\beta^+$ expressing osteoclasts amongst SCI-NHO and TBI-NHO biopsies. (B) Representative immunohistochemistry staining of serial sections of a human NHO biopsies showing the expression of the macrophage marker CD68 (top panel) and the IL-1β protein (bottom panel). Black arrows show IL-1β expressing $CD68^+$ macrophages and red arrows show IL-1β expressing multinucleated $CD68^+$ osteoclasts. 20X magnification and scale bar = 50μm.

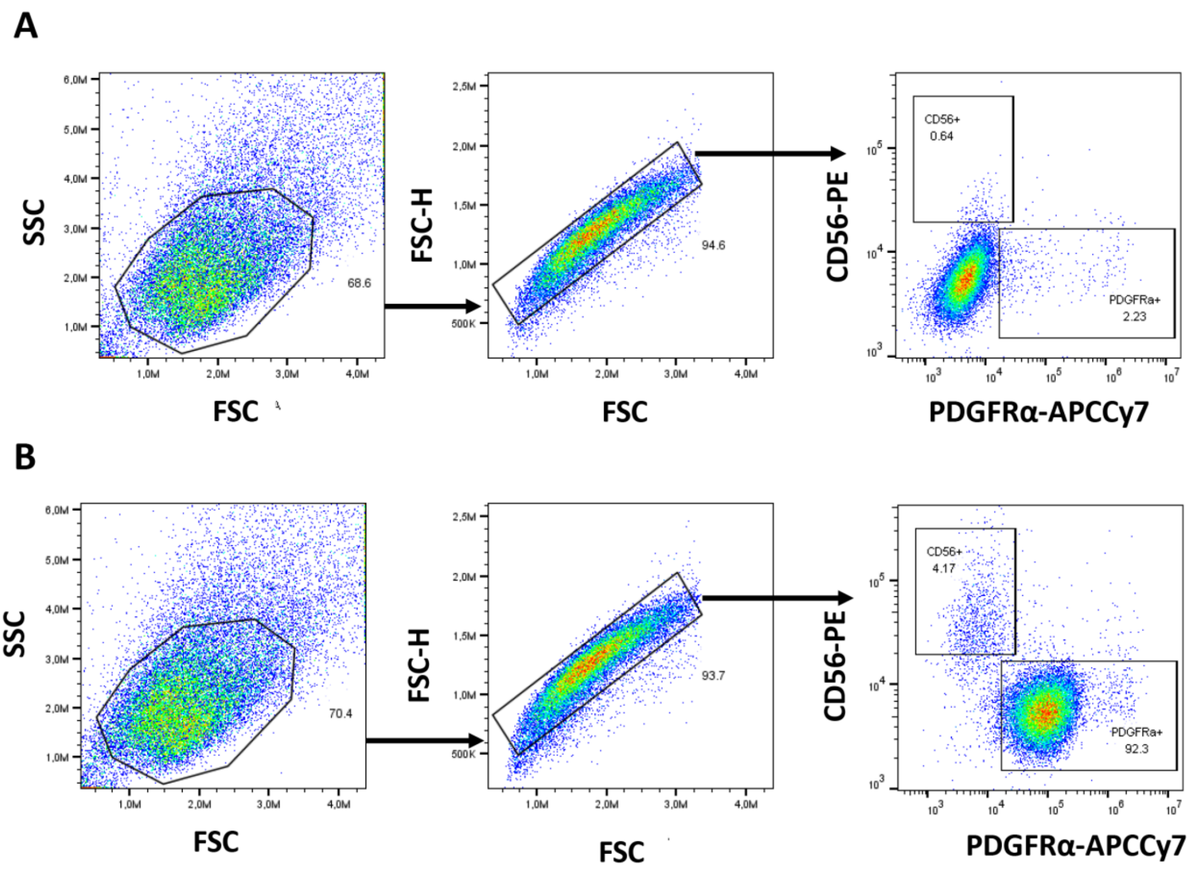


**Supplementary Fig 6.** Flow cytometry gating strategy to isolate human $CD56^{-}PDGFR\alpha^{+}$ FAPs subpopulation from muscle surrounding NHO. From respective singlets gated population (A) Control PE and APC-Cy7 isotype (B) $CD56^{+}PDGFR\alpha^{-}$ satellite cells (SCs) and $CD56^{-}PDGFR\alpha^{+}$ FAPs populations.

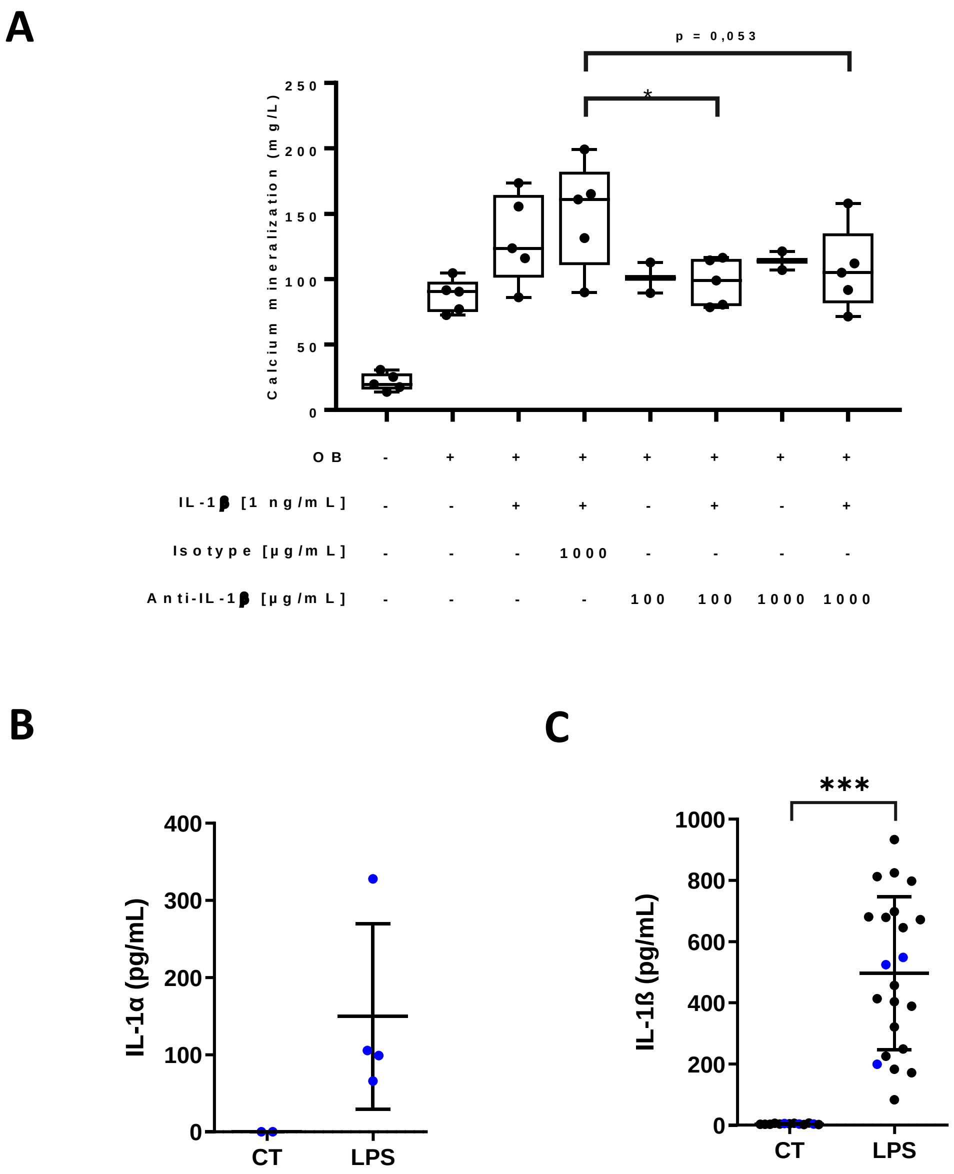


**Supplementary Fig 7.** *In vitro* IL-1β neutralizing strategy during human FAPs osteoblastic differentiation and characterization of human CD14$^+$ derived conditioned medium. (A) Quantification of calcium mineralization of human PDGFRα$^+$ CD56$^-$ FAPs derived from 5 different NHO biopsy / patient cultured with recombinant IL-1β (1 ng/mL) and incubated with anti-IL-1β neutralizing antibody or isotype control as indicated. Each dot represents a result form a different NHO biopsy / patient. Data are represented as boxes extend from the 25th to 75th percentiles, middle line and whiskers represent respectively median and minimum and maximum value. Statistical difference was analyzed using non-parametric repeated measures Friedman test with Dunn's correction for multiple comparisons. *, $p < 0.05$ (B) Concentrations of IL-1α and (C) IL-1β measured by ELISA assays in conditioned media of CD14$^+$ monocytes isolated from NHO bone marrow (black) or from whole blood donation of healthy donors (blue), with no stimulation (CM$^-$) or with LPS stimulation (100 ng/mL, CM$^+$). Data are means ± SD and statistical difference was analyzed using non-parametric two-sided Mann-Whitney test. ***, $p < 0.001$.

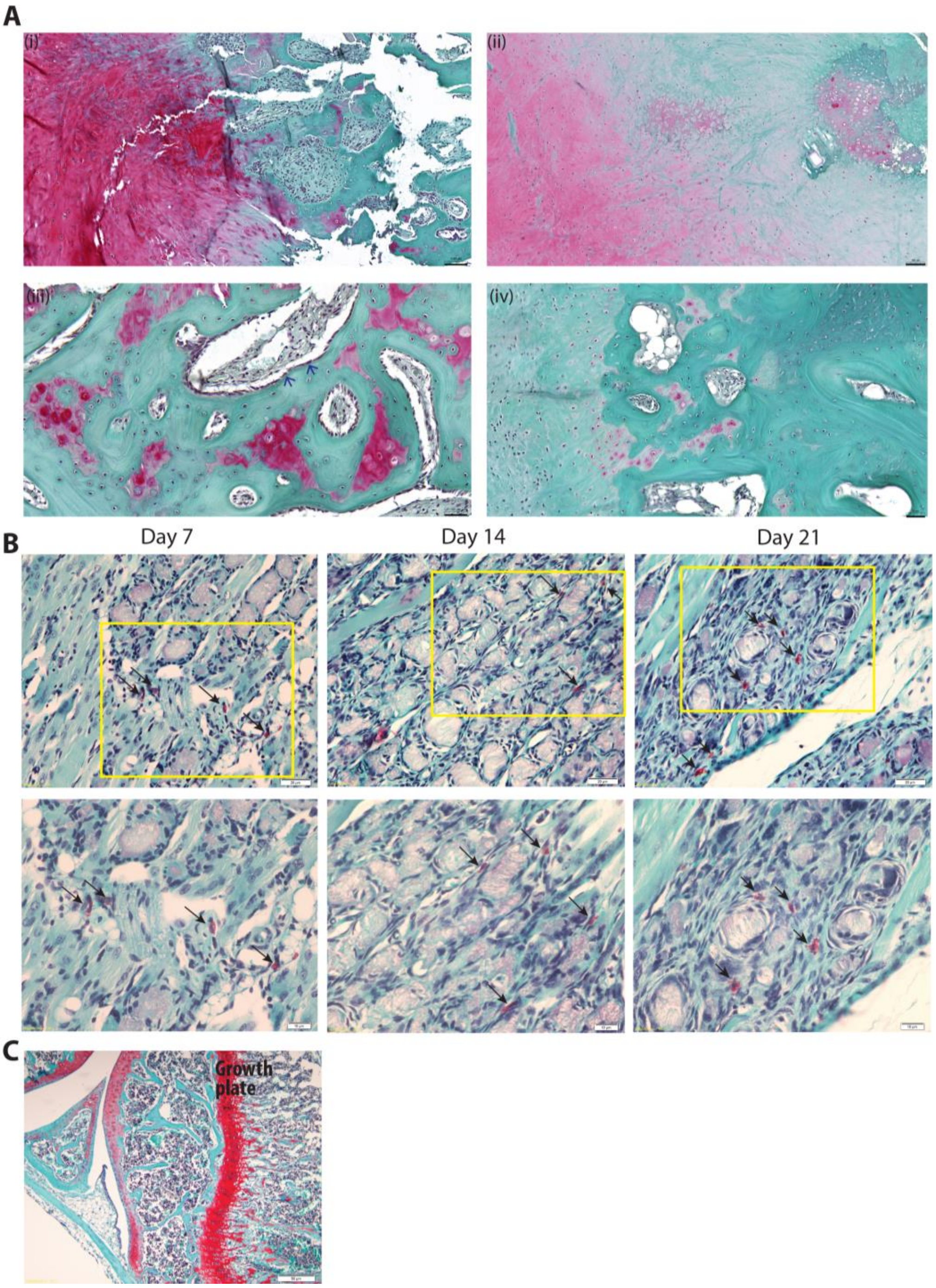


**Supplementary Fig 8.** Endochondral ossification contributes to human NHO formation but not to SCI-NHO mouse model. Safranin O/fast green staining was used to detect the presence of chondrocytes and cartilage matrix, which is stained bright red while bone tissue stained green. (A) (i-iv) Biopsies collected from four NHO patients showed various extent of safranin O staining. (i-ii) Large area with cartilage matrix and chondrocyte-like cells in NHO. (iii) Islands of cartilage were left from remodeling in NHO. (iv) NHO contained very few safranin O positive cells. (B) Images taken from injured muscle of mice underwent SCI+CDTX. No safranin O positive chondrocytes/cartilage were observed in the injured muscle 7, 14 and 21 days post SCI+CDTX surgery (n=3-6 mice per time point). The only safranin O positive cells in injured muscle are mast cells (arrows), characterized by granules that contain proteoglycans. (C) Positive staining in growth plate and articular cartilage shown as positive control of Safranin O staining. Figures in the bottom panel were higher magnification of yellow boxes in the top panel. Scale bars: Panels A(i-ii) 100 µm, A(iii-iv) 50 µm. B top row =20µm, bottom row =10µm. C = 50 µm.

**Supplementary Table 1**: Reagents for qRT-PCR

| | Catalogue number | Company |
|---|---|---|
| TaqMan™ Universal PCR Master Mix | 4323018 | *Thermo Fisher* Scientific |
| *SensiFast* | *BIO65054* | *Bioline* |
| il1b | Mm00434228_m1 | *Thermo Fisher* Scientific |
| tnf | Mm00443258_m1 | *Thermo Fisher* Scientific |
| Osm | Mm01193966_m1 | *Thermo Fisher* Scientific |
| Ccl2 | Mm00441242_m1 | *Thermo Fisher* Scientific |
| Csf1 | Mm00432686_m1 | *Thermo Fisher* Scientific |
| Rps20 | Mm02342828_g1 | *Thermo Fisher* Scientific |

**Supplementary Table 2**. Antibodies for flow cytometry

| Antibody | Catalogue number | Company | Clone | Dilution |
|---|---|---|---|---|
| Anti-B220-FITC | 103206 | Biolegend | RA3-6B2 | 1/200 |
| Anti-Ter119-FITC | 116206 | Biolegend | TER-119 | 1/200 |
| CD3ε-FITC | 100306 | Biolegend | 145-2C11 | 1/300 |
| Anti-LY6G-PECy7 | 127618 | Biolegend | 1A8 | 1/200 |
| CD48 APC-CY7 | 103432 | Biolegend | HM48-1 | 1/200 |
| CD11b-BV510 | 101263 | Biolegend | M1/70 | 1/200 |
| CD45-BV785 | 103149 | Biolegend | 30-F11 | 1/300 |
| Anti-CCR2-PE | FAB5538P | R&D system | 475301 | 1/200 |
| CD11b-FITC | 101206 | Biolegend | M1/70 | 1/100 |
| CD169-PE | 142404 | Biolegend | 3D6.112 | 1/150 |
| Anti-Ly6C-Pacific blue | 128014 | Biolegend | HK1.4 | 1/300 |
| Anti-F4/80-APC | 123116 | Biolegend | BM8 | 1/50 |
| Anti-VCAM1-PECy7 | 105720 | Biolegend | 429 | 1/150 |
| Fixable Viability Stain 700 | 564997 | BD Bioscience | - | 1:15000 |
| 7AAD | A1310 | Life Technologies | - | |
